# Systematic identification of structure-specific protein–protein interactions

**DOI:** 10.1101/2023.02.01.522707

**Authors:** Aleš Holfeld, Dina Schuster, Fabian Sesterhenn, Patrick Stalder, Walther Haenseler, Inigo Barrio-Hernandez, Dhiman Ghosh, Jane Vowles, Sally A. Cowley, Luise Nagel, Basavraj Khanppnavar, Pedro Beltrao, Volodymyr M. Korkhov, Roland Riek, Natalie de Souza, Paola Picotti

## Abstract

Protein–protein interactions (PPIs) mediate numerous essential functions and regulatory events in living organisms. The physical interactome of a protein can be abnormally altered in response to external and internal cues, thus modulating cell physiology and contributing to human disease. In particular, neurodegenerative diseases due to the accumulation of aberrantly folded and aggregated proteins may lead to alterations in protein interactomes. Identifying changes in the interactomes of normal and disease states of proteins could help to understand molecular disease mechanisms, but current interactomics methods are limited in the ability to pinpoint structure-specific PPIs and their interaction interfaces on a proteome-wide scale. Here, we adapted limited proteolysis–mass spectrometry (LiP–MS) to systematically identify putative structure-specific PPIs by probing protein structural alterations within cellular extracts upon treatment with specific structural states of a given protein. We demonstrate the feasibility of our method to detect well-characterized PPIs, including antibody–target protein interactions and interactions with membrane proteins, and show that it pinpoints PPI interfaces. We then applied the LiP–MS approach to study the structure-specific interactors of the Parkinson’s disease hallmark protein alpha-synuclein (aSyn). We identified several previously known interactors of both aSyn monomer and amyloid fibrils and provide a resource of novel putative structure-specific interactors for further studies. This approach is applicable to identify structure-specific interactomes of any protein, including posttranslationally modified and unmodified, or metabolite-bound and unbound structural states of proteins.

## Introduction

Many cellular processes are governed by proteins assembled into complexes; thus, protein–protein interactions (PPIs) have multiple essential roles in cells. The physical interactome of a given protein (*i.e*., the set of proteins with which it interacts) is not static. The organization of the interactome can be altered due to numerous molecular events that occur in response to environmental stimuli, stress, time, and disease state (Goh *et al*, 2007). These molecular events include not only genetic variations (Carter *et al*, 2013; Ferlini & Fini, 2015; Auton *et al*, 2015) but also covalent (Mann & Jensen, 2003; Pan & Chen, 2022; Xu *et al*, 2018; Jensen, 2004; Khoury *et al*, 2011; Duan & Walther, 2015) and noncovalent modifications (Schmidt & Robinson, 2014; Gillingham *et al*, 2019) that can lead to structural alterations of a given protein. Thus, a protein of interest may associate with different sets of protein partners under normal compared to disease conditions.

Abnormal alterations in PPIs have the potential to modulate physiological processes and contribute to disease phenotypes (Sahni *et al*, 2015; Thompson *et al*, 2020; Calabrese *et al*, 2022). For example, in neurodegenerative diseases such as Parkinson’s disease (PD), dementia with Lewy bodies (DLB), multiple system atrophy (MSA), Alzheimer’s disease (AD), and Huntington’s disease, disease-associated proteins (*e.g*., alpha-synuclein (aSyn), amyloid-β, tau or huntingtin) aggregate into β-sheet-rich structures that are thought to be toxic to cells (Soto, 2003; Goedert, 2015; Bates, 2003; Taylor *et al*, 2002; Ross & Poirier, 2004). However, it remains enigmatic how protein aggregation affects cell physiology. One hypothesis is that aggregation-prone proteins, such as aSyn, may undergo changes in their interactomes while transitioning from the monomeric to the aggregated state (Leitão *et al*, 2021; van Diggelen *et al*, 2020; Lassen *et al*, 2016; Betzer *et al*, 2015). Such interactome changes could profoundly affect cellular physiology and could play a role in the onset of various diseases. Thus, a systematic assessment of structure-specific interactomes could help elucidate pathological cellular processes and unravel disease mechanisms.

Multiple methods have been developed to study PPIs (Meyerkord & Fu, 2015), but all have limitations for the study of structure-specific interactomes. Affinity purification coupled to mass spectrometry (AP–MS) relies on purification of a bait protein of interest from a cellular extract, together with its interaction partners (Dunham *et al*, 2012; Collins & Choudhary, 2008; Morris *et al*, 2014; Meyer & Selbach, 2015; Chang, 2006). These experiments typically only detect stable interactions, and engineered affinity tags may alter protein structures and interaction sites. Furthermore, structural changes of proteins may alter interactions with specific antibodies thus affecting the capability to detect structure-specific interactomes, and structure-specific antibodies are not available for most proteins (Kumar *et al*, 2020). Proximity labeling approaches, such as BioID (Roux *et al*, 2012) or APEX (Martell *et al*, 2012), identify interacting proteins by fusing one or more baits with an enzyme that covalently labels proximal proteins (Go *et al*, 2021; Trinkle-Mulcahy & Poterszman, 2019; Han *et al*, 2018; Xu *et al*, 2021; Samavarchi-Tehrani *et al*, 2020). Although these strategies allow the capture of more transient interactions and can be employed in living cells, they also identify bystander proteins that are near the bait but do not interact with it. The interactome can also be profiled in an untargeted manner using co-fractionation techniques coupled to MS (Kirkwood *et al*, 2013; Bludau *et al*, 2021; Heusel *et al*, 2019; Bludau *et al*, 2020; Heusel *et al*, 2020), in which proteins are separated according to size/shape (size exclusion chromatography; SEC) or charge (ion exchange chromatography), and interactions are inferred based on co-fractionation patterns (Scott *et al*, 2015; Hu *et al*, 2019; Fossati *et al*, 2021; Stacey *et al*, 2017). SEC–MS has identified thousands of putative PPIs (Heusel et al, 2019; Bludau et al, 2021, 2020; Heusel et al, 2020; Kristensen et al, 2012; Liu et al, 2008), examined protein complex dynamics (Kristensen *et al*, 2012), and improved the detection of variations in protein complexes associated with specific proteoforms (Kirkwood *et al*, 2013). However, these methods are not easily scalable and do not report directly on physical interactions, which can lead to false positive assignments. Furthermore, different structural states of proteins may be insufficiently separated in the chromatographic step, and studying PPIs of aggregated proteins can be hindered by the elution of aggregates in the void volume together with both interacting and non-interacting proteins. Finally, in crosslinking coupled to MS (XL–MS) (Iacobucci *et al*, 2020; Wheat *et al*, 2021; Leitner *et al*, 2010; Liu *et al*, 2017; Holding, 2015; Leitner *et al*, 2020; Chavez & Bruce, 2019; Liu & Heck, 2015; Leitner *et al*, 2016), covalent links are formed between proximal amino acid residues to probe PPIs as well as three-dimensional structures and interaction interfaces; however, due to the difficulty of identifying crosslinked peptides, XL–MS yields only modest coverage of the interactome in complex biological samples.

In this study, we report an approach for the detection of PPIs in complex proteomes based on limited proteolysis– mass spectrometry (LiP–MS) (Schopper *et al*, 2017; Feng *et al*, 2014; Malinovska *et al*, 2022), our previously developed structural proteomics method that relies on the brief application of a sequence-unspecific protease, proteinase K, to a cellular extract under native conditions followed by trypsin digestion. These steps generate structure-specific proteolytic fragments that can be measured with MS. We have previously shown that LiP–MS detects protein structural changes (Feng *et al*, 2014), metabolite– and drug–protein interactions (Holfeld *et al*, 2023; Piazza *et al*, 2018, 2020), and other functional events within complex cellular extracts with peptide-level resolution (Cappelletti *et al*, 2021). We postulated that LiP–MS would detect PPIs since physical interaction between two proteins should alter their protease accessibility either at the interaction interface itself or in other protein regions that change structurally upon interaction (Konno, 1987; Wilson, 1991; de Pereda & Andreu, 1996; Digiacomo *et al*, 2017). Thus, adding a protein to a cellular extract should result in altered protease accessibility of its cellular interactors. These changes in proteolytic patterns could then be detected by quantitative MS analysis to identify interactors of the target protein. Importantly, adding distinct structural states of a protein to the cell extract should enable comparison of their interactomes and thus identification of structure-specific interactions. Here, we demonstrate that LiP–MS can be applied to the systematic investigation of PPIs in complex cellular environments and to detect structure-specific interactomes. We show that the approach detects known interactions between the respiratory syncytial virus F glycoprotein and its site-specific antibodies, including the identification of several known antigenic sites, directly in a eukaryotic cellular environment. Therefore, the approach enables the identification of protein–protein interaction interfaces and may estimate relative binding parameters. The method can also be applied to study PPIs of integral membrane proteins, which we demonstrate on the interaction between adenylyl cyclase 8 and calmodulin, as proof of principle. Finally, we applied LiP–MS to study structure-specific interactomes of aSyn, a protein involved in PD, for which the mechanisms of toxicity are still largely unknown. In summary, the detection of structure-specific interactors of disease-associated protein structural states should provide novel molecular insights into disease mechanisms and suggest new therapeutic targets.

## Results

### Conformation-specific protein–protein interactions detected by LiP–MS

We tested the feasibility of identifying PPIs using the LiP–MS workflow (Figure 1a) by probing well-characterized interactions *in vitro*. First, we investigated interactions between respiratory syncytial virus F (RSVF) glycoprotein and several site-specific neutralizing monoclonal antibodies against this target. The RSVF glycoprotein is a class I fusion protein that undergoes a conformational change from a metastable prefusion state to a stable postfusion state during viral entry. We incubated the purified recombinant RSVF glycoprotein stabilized in its prefusion or postfusion state with each of five purified antibodies specific for one of the three antigenic sites Ø, II, or IV (Figure 1b) or with an unspecific human IgG antibody (referred to as control). We applied the LiP–MS workflow and identified antibody-induced structural alterations of RSVF based on LiP peptide intensities that were significantly different (log_2_-fold change, |log_2_ FC| > 1; *q*-value < 0.01, moderated t-test) between a sample incubated with each site-specific antibody versus control; MS analysis was performed using label-free data-independent acquisition (DIA). We then mapped the significantly altered peptides in preRSVF (PDB: 4JHW) (McLellan *et al*, 2013) and postRSVF (PDB: 3RRR) (McLellan *et al*, 2011) onto the three-dimensional structures of the relevant protein conformation (Figure 1c).

**Figure 1:**
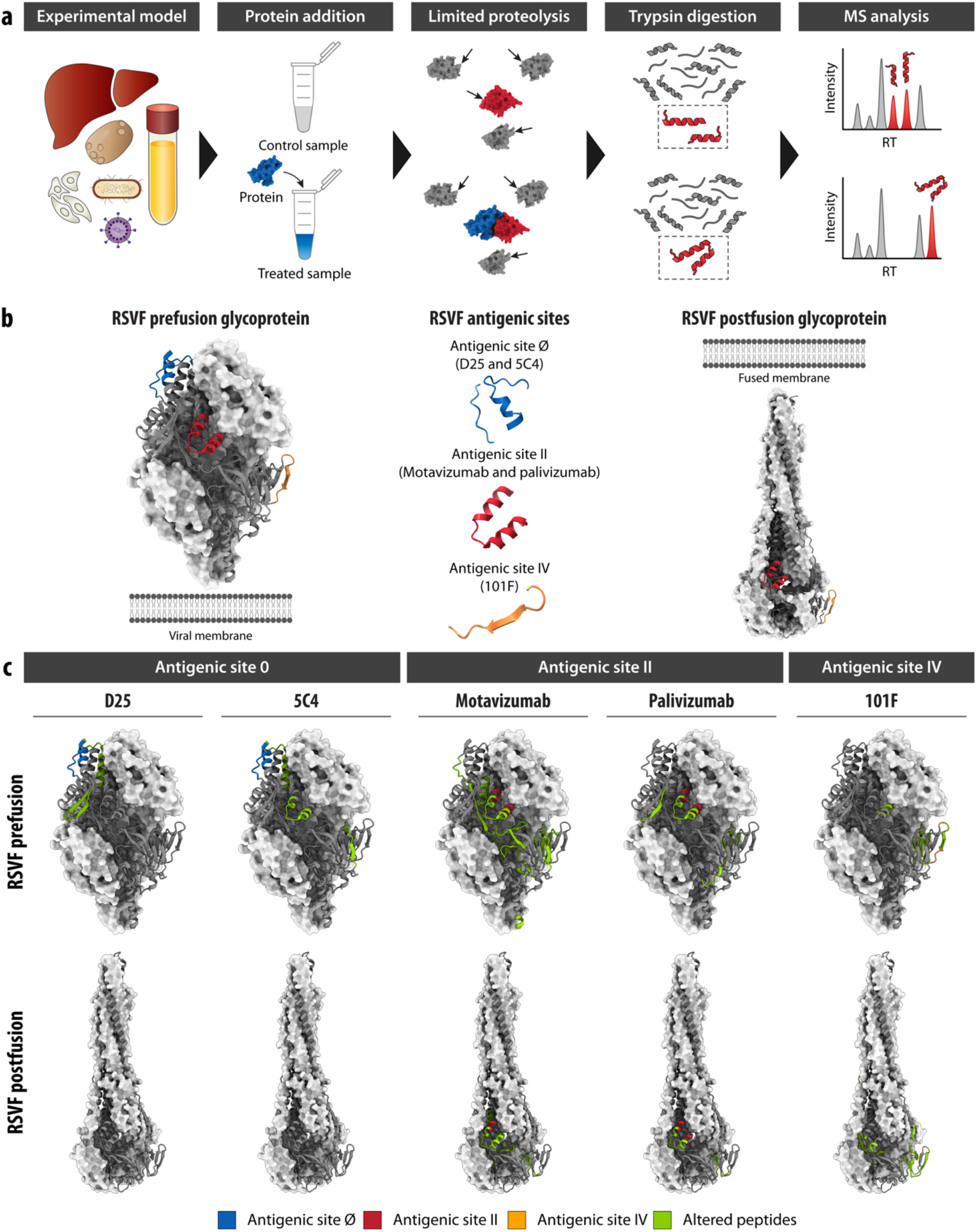
LiP–MS detects protein–protein interactions in purified systems. (a) Schematic of LiP–MS workflow. Proteins are extracted from an experimental model, such as tissues, human cells, bacteria, yeast, viruses, or biofluids, under native-like conditions. The extract is then exposed to a protein of interest (treated) or not exposed (control) and subjected to limited proteolysis with proteinase K. Under LiP conditions, proteinase K cleaves solvent-exposed, accessible, and flexible regions thus generating protein fragments that may differ between the treated and control samples for an interactor of the spiked-in protein. These protein fragments are digested by trypsin under denaturing conditions to produce peptides that are measurable by bottom-up proteomics. By comparing differential peptides between the treated and control sample, interactors of the protein of interest can be identified. (b) Structures of preRSVF (left, PDB: 4JHW) (McLellan *et al*, 2013) and postRSVF (right, PDB: 3RRR) (McLellan *et al*, 2011). Known antigenic sites are shown both on the protein structure and in isolation (middle). Blue indicates antigenic site Ø, targeted by antibodies D25 and 5C4. Red indicates antigenic site II, targeted by palivizumab and motavizumab. Orange indicates antigenic site IV, targeted by 101F. (c) Visualization of structurally altered peptides (|log_2_ FC| > 1, moderated t-test, *q*-value < 0.01) in green, on one of the subunit of trimeric preRSVF (upper panel) and postRSVF (lower panel) protein structures upon addition of the indicated antibodies. Antigenic sites are colored as in panel (b).

Antigenic site Ø is situated at the trimer apex of preRSVF and consists of a kinked α helix (17 residues) and a disordered loop (7 residues) (McLellan *et al*, 2013). Antibodies targeting this site (D25 and 5C4) are known to be specific for the prefusion conformation of the RSVF glycoprotein. Consistent with this, incubation with the D25 antibody resulted in four significantly changed LiP peptides on the preRSVF protein (out of 412 detected peptides) (Supplementary Data 1). No significant changes were observed for peptides of the postfusion protein (out of 427 detected peptides). Furthermore, one of the four altered peptides in the prefusion protein mapped directly to the antigenic site Ø, and the other three altered peptides were situated near the antigenic site Ø (Figure 1c; we define direct mapping as the identified peptide containing the sequence of the known antigenic site), confirming the binding of D25 to the expected RSVF region. Similarly, the addition of the 5C4 antibody yielded seven significantly changed peptides on the prefusion form of RSVF (out of 447 detected peptides), one of which mapped to the antigenic site Ø, whereas we detected no significant changes for the postfusion protein (out of 408 detected peptides). Although both 5C4 and D25 bind to the antigenic site Ø, these antibodies are known to differ in their vertical and horizontal angles of approach (Tian *et al*, 2017), which may explain the detection of altered LiP peptides at the antigenic site II.

The highly conserved antigenic site II is found on both prefusion and postfusion conformations of RSVF glycoprotein and is recognized by the antibodies palivizumab (Synagis®) and motavizumab (MEDI-524, Numax). For the RSVF prefusion glycoprotein, we identified eight peptides with altered abundances relative to the control due to palivizumab and 31 due to motavizumab binding (out of 453 and 412 detected peptides, respectively) (Supplementary Data 1). Seven of these peptides showed changes for both antibodies and mapped at or near the known antigenic site II. For the postfusion conformation, we detected five significantly changed peptides upon palivizumab binding (out of 432 detected peptides), four of which were also detected for preRSVF and which again mapped directly to the antigenic site II. For motavizumab, we found 12 differential peptides (out of 414 detected peptides) compared to the control, which were likewise situated either directly at or near the antigenic site II (Figure 1c). The greater number of significantly altered peptides for motavizumab could be because it is an enhanced potency antibody developed from palivizumab and, as such, binds to the target protein with much higher affinity. This is supported by our finding that relative abundance changes of altered RSVF peptides were larger when motavizumab was bound. This observation is also in good agreement with the recent report that small molecules that bind with higher affinity result in higher occupancy and thus larger fold changes in LiP peptide abundances (Piazza *et al*, 2018, 2020).

Antigenic site IV on the RSVF glycoprotein involves an irregular six-residue bulged β-strand epitope and is the major target of 101F antibody in both prefusion and postfusion forms (McLellan *et al*, 2010). We observed eight peptides (out of 426 detected peptides) on preRSVF and 11 peptides (out of 428 detected peptides) on postRSVF that changed proteolytic patterns upon 101F binding (Supplementary Data 1). As expected, all peptides mapped either at or near antigenic site IV (Figure 1c). In summary, our data show that LiP–MS detects target protein-antibody interaction interfaces for several well-characterized target protein-antibody pairs under defined, purified conditions. These findings support our hypothesis that LiP–MS can be used to pinpoint protein regions that mediate interactions between an antibody and its target protein, including conformation-specific interactions.

Since our goal was to systematically identify PPIs within a native cellular environment, we further analyzed the interactions between postRSVF and motavizumab in a complex extract of HEK293T cells. We identified 14 peptides (|log_2_ FC| > 0.75; moderated t-test, *q*-value < 0.01; Supplementary Data 2) that significantly changed in LiP intensity upon addition of 3 μg motavizumab to the lysate, corresponding to seven proteins (out of 69,263 detected peptides, corresponding to 4,582 proteins). Of the 14 changing peptides, eight mapped to the postRSVF glycoprotein, with 2 out of 8 peptides overlapping with peptides detected *in vitro*. Structural changes detected in the other six proteins could have resulted from direct interactions of RSVF or motavizumab with these proteins in the lysate or from indirect structural effects. Similarly, contaminant proteins present in the preparations of postRSVF or the antibody could interact with proteins in the lysate or otherwise cause indirect structural changes. Next, we performed a dose-response experiment to better distinguish between true and false positive hits, as we have previously done to identify small-molecule–protein interactions within complex proteomes using LiP–MS (Piazza *et al*, 2020; Holfeld *et al*, 2023). We exposed the HEK293T cellular extract, supplemented with 1 μg of postRSVF, to five concentrations of motavizumab, and identified peptides that showed high correlation (*r*) to a sigmoidal trend of the peptide-intensity response profile. Of the 14 peptides identified in the single-dose experiment, the intensity responses of eight peptides were proportional to the amount of motavizumab with high correlation (*r* > 0.85; Figure 2a). Importantly, all eight were postRSVF peptides, and all were mapped onto or near the antigenic site II (Figure 2b, c). In summary, *in situ* LiP–MS analysis enables an unbiased identification of PPIs and pinpoints interaction interfaces in complex biological matrices.

**Figure 2:**
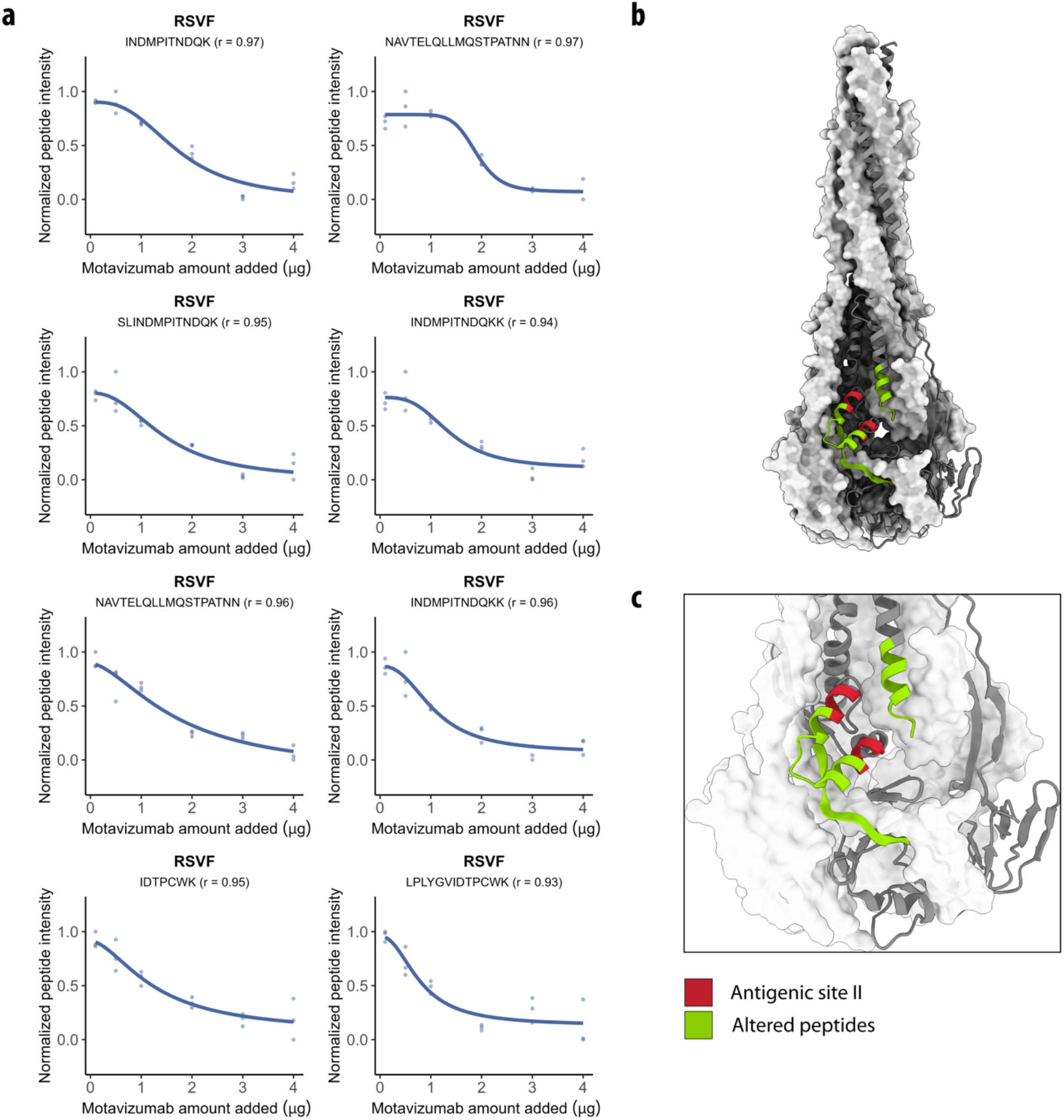
LiP–MS detects protein–protein interactions in complex proteomes. (a) Dose-response curves of eight LiP peptides originating from postRSVF show relative peptide intensities proportional to the amount of motavizumab spiked into HEK293T cellular extracts. Pearson’s coefficient (r) to a sigmoidal trend of the peptide-intensity response profile is indicated. (b) The structure of postRSVF (PDB: 3RRR) (McLellan *et al*, 2011) with peptides altered in the dose-response analysis (*r* > 0.85; |log_2_ FC| > 0.75; moderated t-test, *q*-value < 0.01) indicated in green and antigenic site II in red. (c) Zoom of the altered peptides on the structure of postRSVF (PDB: 3RRR) (McLellan *et al*, 2011) with colors as in panel (b).

### LiP–MS detects protein–protein interactions with integral membrane proteins

Integral membrane proteins (IMPs) represent a biologically interesting set of proteins as they constitute a large proportion of therapeutic targets in drug discovery. However, IMPs and their interacting proteins remain challenging to measure in bottom-up and structural proteomics experiments. Therefore, we asked whether LiP– MS enables the identification of PPIs of membrane proteins. We tested the applicability of LiP–MS to detect the interaction of calmodulin (CaM) with membrane-integral adenylyl cyclase type 8 (AC8). CaM is an intracellular Ca^2+^-binding protein that is known to interact with CaM-binding domains (CaMBDs) in the N-terminus and in the C-terminal cytoplasmic regulatory subdomain (AC8-C2b) of AC8 (Figure 3a) (Gu & Cooper, 1999; Herbst *et al*, 2013). We applied the LiP–MS workflow to crude membrane preparations from HEK293F GnTI- cells engineered to overexpress bovine AC8 fused at its C terminus to TwinStrep-YFP, incubated with a 6-dose concentration series of bovine CaM. The coverage of membrane-annotated proteins was better in the crude membrane preparations than in standard cell lysates from which membranes had been removed (Figure 3b). We also observed good sequence coverage of our target AC8 (220 peptides covering 58.5% of the AC8 sequence) in the crude membrane preparation, although we did not detect peptides from the transmembrane domains (Figure 3c), as expected in any bottom-up proteomics experiment, due to their hydrophobicity.

**Figure 3:**
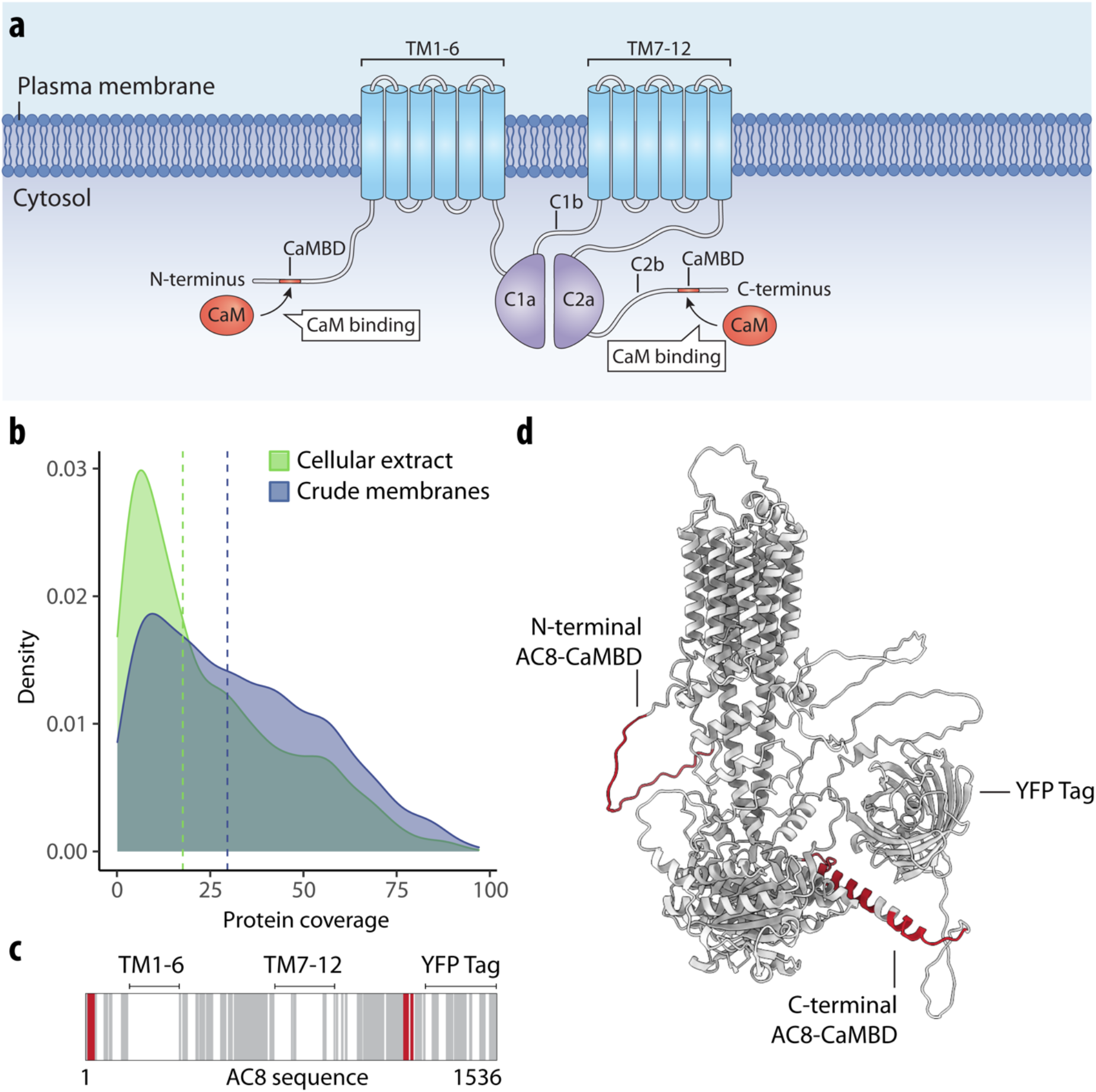
LiP–MS detects interactors of integral membrane proteins in crude membranes. (a) Schematic of AC8 with the CaMBD in the N-terminus, transmembrane domains 1-6 and 7-12 (TM1-6 and TM7-12), and catalytic domains C1a, C1b, C2a, and C2b indicated. (b) Distribution of protein coverage for membrane-annotated proteins identified in crude membrane preparations of HEK293F GnTI- cells (blue) and in HEK293T cellular extracts (green). Blue and green vertical lines indicate calculated median coverages of 29.6% and 17.6%, respectively. (c) Protein sequence coverage of bovine AC8-YFP in LiP–MS is visualized using a barcode, which depicts peptides detected along the AC8-YFP sequence. Gray represents detected peptides, white represents non-detected regions, and red represents peptides that were significantly altered upon CaM addition (*r* > 0.85, |log_2_ FC| > 1, moderated t-test, *q*-value < 0.01). (d) AlphaFold2 (Varadi *et al*, 2022; Jumper *et al*, 2021) predicted the structure of AC8 (including the tag domain) with peptides altered upon CaM addition highlighted in red.

Upon addition of CaM to the membrane preparations, 279 peptides were significantly altered (of 91,847 peptides detected, corresponding to 5,185 proteins) relative to the no-treatment control (*r* > 0.85, |log_2_ FC| > 0.75, moderated t-test, *q*-value < 0.01; Supplementary Data 3). These peptides mapped to 163 proteins. Amongst these, 16 peptides with high correlation to sigmoidal profiles (*r* > 0.85; Figure S 1) originated from AC8 and mapped exactly to the N-terminal AC8-CaMBD and the C-terminal AC8-CaMBD. These data confirmed that LiP–MS detects CaM binding and pinpoints known binding sites.

We then examined the larger set of proteins that underwent structural alterations upon CaM addition to the crude membrane preparation. We searched for canonical CaM-binding motifs within the sequences of all proteins for which we detected structural alterations upon CaM addition and showed that 85 of the 279 significantly altered peptides (corresponding to 56 proteins) are predicted to contain CaM-binding motifs (Mruk *et al*, 2014). Overall, our data demonstrate that the LiP–MS pipeline detects protein interactors of IMPs and soluble proteins and enables the identification of interaction interfaces *in situ* in detergent-free crude membranes.

### Differential interactomes of alpha-synuclein monomer and amyloid fibrils

Parkinsons’s disease (PD) is associated with the aggregation of the protein alpha-synuclein (aSyn) into fibrillar structures in neuronal cells, but mechanisms of aSyn toxicity remain unclear (Wong & Krainc, 2017). One current hypothesis is that, upon aggregation, aSyn undergoes changes in its interactome that underlie disease development (Leitão *et al*, 2021; Lassen *et al*, 2016; Betzer *et al*, 2015; van Diggelen *et al*, 2020). We thus applied our LiP– MS approach to assess whether monomeric and aggregated, fibrillar structural states of aSyn have different cellular interactomes. We generated a cellular extract of cortical neurons differentiated from an SNCA-knockout (KO) induced pluripotent stem cell (iPSC) line (Figure S 2), to avoid possible effects of endogenous aSyn on the analysis (Fernandes *et al*, 2016; Zambon *et al*, 2019; Haenseler *et al*, 2017a). We purified acetylated aSyn monomer, which is considered to be the physiologically relevant form (Burré *et al*, 2013; Fauvet *et al*, 2012; Runfola *et al*, 2020), and generated aSyn amyloid fibrils *in vitro*, ensuring the conformations of our preparations using SEC, TEM, native-PAGE, and ThT fluorescence as quality control steps (Figure S 3). Subsequently, we spiked increasing amounts of aSyn monomer or fibrils into lysates of SNCA-KO iPSC-derived neurons. We then performed LiP–MS experiments in a dose-dependent manner to identify the resulting protein structural alterations across the proteome and thus putative interactors of the monomeric and amyloid fibril conformational states of the protein.

We identified 68 and 242 significantly changing peptides upon addition of aSyn monomer and amyloid fibrils to the cellular extracts, respectively (*r* > 0.85, |log_2_ FC| > 0.75, moderated t-test, *q*-value < 0.01; Supplementary Data 4) (out of 90,416 and 85,084 detected peptides, corresponding to 5,435 and 5,536 proteins) (Figure 4a). A total of 64 proteins showed structural changes upon spike-in of aSyn monomer and 225 proteins upon spike-in of aSyn fibrils, indicating a higher apparent binding to aSyn fibrils compared to monomer. Several putative aSyn interacting proteins displayed monomer-specific (n = 50; Supplementary Data 4) and fibril-specific changes (n = 211; Supplementary Data 4) (Figure 4b). In general, 14 putative interactors were found to be conformation-unspecific (CANX, CCT2, EEF1A1, FARSB, PAF1, PEBP1, PIN1, RBM8A, RPS27A, SEC13, SMC3, SPTAN1, VPS52, and YWHAB), nine of which are known vesicular proteins, thus supporting evidence that aSyn localizes with vesicles (Ebanks *et al*, 2020). Five of the 14 proteins were previously reported to bind aSyn in the STRING database (CANX, EEF1A1, PAF1, PIN1, RPS27A).

**Figure 4:**
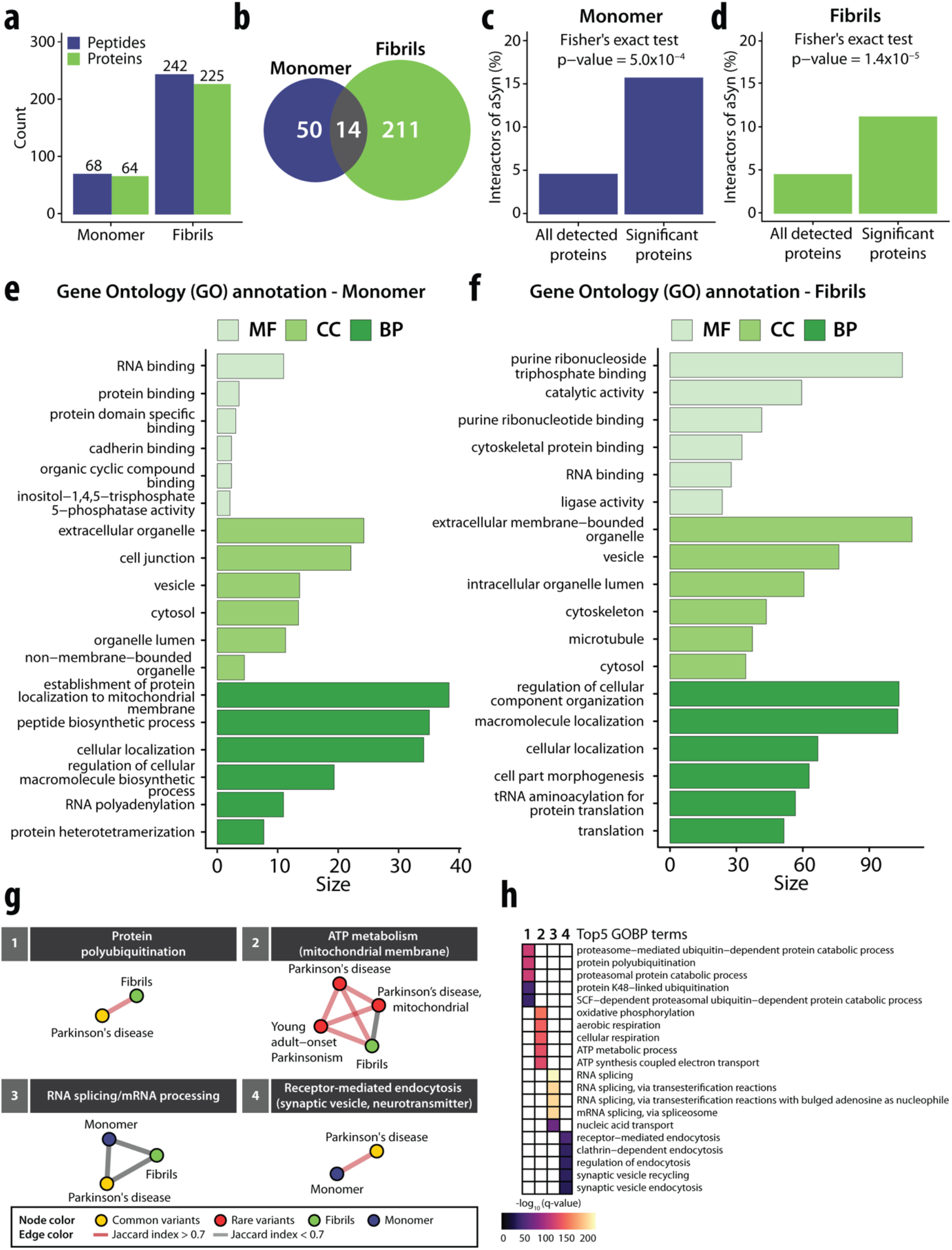
A systematic investigation of structure-specific interactors of the amyloidogenic protein aSyn and gene module analysis of Parkinson’s disease. (a) Barplot with the numbers of altered LiP peptides (blue) and corresponding structurally altered proteins (green) for aSyn monomer (left) and amyloid fibrils (right). (b) Venn diagram with the numbers of structurally altered proteins for aSyn monomer (blue), and for amyloid fibrils (green). The overlap of structurally altered proteins identified for both aSyn monomer and amyloid fibrils is indicated in grey. (b) The plots show the fraction of known aSyn interactors (based on the STRING database) in structurally altered proteins (right) versus all detected proteins (left) upon spike-in of aSyn monomer into an iPSC-derived cortical neuron extract. The *p*-value assessing enrichment (Fisher’s exact test) is shown. (d) Enrichment plot as in (c) upon spike-in of aSyn fibrils. (e-f) Functional enrichment analyses of structurally altered proteins upon spike-in of aSyn monomer (e) or fibril (f), based on the indicated ontologies (molecular function in light green, cellular component in green, biological process in dark green); the plots show the size (i.e., a score calculated based on *q*-value) of the top6 significant gene ontology terms upon removal of redundant terms (*q*-value < 0.01, Benjamini-Hochberg FDR, minimum hypergeometric test, SimRel functional similarity, size = 0.7). (g) Identified modules with enriched GOBP (*q*-value < 0.05, Fisher’s exact test, one-sided) that are linked to either common (yellow node) or rare variants (red node) of PD genes for aSyn monomer (blue) or fibrils (green). The thickness of lines represents the Jaccard index (red for Jaccard index > 0.7, gray for Jaccard index < 0.7). (h) Heatmap showing the top5 GOBP terms within each module as indicated in (b). The gradient color indicates the significance based on the results of a GOBP enrichment test (purple = low significance, yellow = high significance).

First, we analyzed proteins (n = 64; Supplementary Data 4) that showed structural changes upon treatment with aSyn monomer. This set of proteins was significantly enriched (*p*-value < 0.01, Fisher’s exact test) for known interactors of aSyn, containing ten proteins that were previously classified as physical interactors of aSyn in the STRING database (CALM1, CANX, EEF1A1, ILF3, MAP1B, PAF1, PIN1, RPS27A, VIM, and YWHAZ) (Figure 4c). Interestingly, the interaction between CALM1 and aSyn was reported to be monomer-specific (Lee *et al*, 2002), consistent with our data. In addition, we identified structural changes in several proteins, such as AGRN, SYNJ1, MAP1B, and YWHAZ, which have been implicated in PD based on disease-gene associations mined from literature, and in peroxiredoxin-1 (PRDX1), which has been linked to neurodegenerative processes (Hallacli *et al*, 2022; Szeliga, 2020). A functional enrichment (GO) analysis of the putative interactors of aSyn monomer showed enrichment for RNA-binding, protein-binding, and protein specific domain-binding molecular functions (*q*-value < 0.01, Benjamini-Hochberg FDR, minimum hypergeometric test; SimRel functional similarity, size = 0.7) (Figure 4e), consistent with the known interaction of aSyn with proteins involved in mRNA translation (Hallacli *et al*, 2022; Chung *et al*, 2017). Putative interactors were also enriched for extracellular organelles, cell junction, and vesicles (GO cellular components), consistent with the known localization of aSyn to presynaptic terminals, its interaction with synaptic vesicles, and its roles in exocytosis, endocytosis, and vesicle recycling (Huang *et al*, 2019). Finally, putative interactors were enriched (GO biological processes) for establishment of protein localization to mitochondrial membrane, peptide biosynthetic process, and cellular localization, supporting prior evidence for aSyn monomer involvement in mitochondrial bioenergetics (Ludtmann *et al*, 2016) and membrane transport (Huang *et al*, 2019).

Next, we examined proteins (n = 225; Supplementary Data 4) that were structurally altered in neuronal lysates upon spike-in of aSyn fibrils. This set was again enriched in known aSyn interactors (n = 25 proteins; CALR, CANX, CCT3, DNMT1, DYNC1H1, DYNLL1, EEF1A1, EFTUD2, FKBP1A, GAPDH, HSP90AA1, HSP90AB1, HSPA1A, HSPA8, HSPD1, MAP2K2, PAF1, PIN1, PPP3CA, PREP, RAB3A, RPS27A, RPS3, SMU1, and SNRNP200) (*p*-value < 0.01, Fisher’s exact test) (Figure 4d). As for monomeric aSyn, we identified proteins previously reported to interact exclusively with aggregated forms of aSyn: the mitochondrially localized protein superoxide dismutase 2 (SOD2) which interacts specifically with fibrillar aSyn, and the vesicular ras-related protein Rab-3A (RAB3A) which preferentially interacts with oligomeric and aggregated aSyn (Chen *et al*, 2013; Tan *et al*, 2022; Huang *et al*, 2018). Among other proteins structurally altered upon addition of aSyn fibrils, we also identified the amyloidogenic protein gelsolin (GSN), a component of PD-associated intraneuronal inclusions of which aggregated aSyn is a major component (Welander *et al*, 2011), as well as the actin-binding protein cofilin-1 (CFL1), which is known to co-aggregate with aSyn fibrils and is implicated in pathogenicity in PD (Tan *et al*, 2022; Yan *et al*, 2022). Notably, we further identified four components of chaperonin-containing T-complex (CCT2, CCT3, CCT6A, and CCT8) that play a central role in protein folding, degradation, aggregation, as well as in the inhibition of aSyn aggregation (Ghozlan *et al*, 2022; Sot *et al*, 2017; Grantham, 2020).

We compared our data with previous proteomic analyses of the content of Lewy Bodies (LBs) either in a neuronal aSyn fibril seeding model (Mahul-Mellier *et al*, 2020) or in postmortem patient brains (Petyuk *et al*, 2021; Xia *et al*, 2008). We found an enrichment of LB-associated proteins in the set of putative aSyn fibril interactors. In total, 49 and 38 of our fibril-dependent structurally altered proteins (out of 225 detected proteins) were previously identified as components of neuronal LBs after 14 and 21 days of cell treatment with aSyn fibrils, respectively, with a significant enrichment over all detected proteins that overlapped with the previous study (734/422 respectively) (*p*-value < 0.01, Fisher’s exact test; Figure S 4a, Figure S 4b). When we compared our data with a previous study that investigated LBs purified from *postmortem* brain tissues of patients diagnosed with the LB variant of AD (Xia *et al*, 2008), we found that 14 of the proteins with structural changes (*p*-values < 0.05 and an absolute fold-change |FC| > 0.5) overlapped with the previously defined LB proteins, of the 67 overlapping proteins detected overall, with a significant enrichment (*p*-value < 0.01, Fisher’s exact test) (Figure S 4c). This set of proteins included six known aSyn interactors (CALR, CANX, DYNC1H1, GAPDH, HSP90AB1, and RPS3), as well as several mitochondrial proteins (e.g., ACO2, ATP5PB, IDH2, NDUFS1, and RPS3), calcium-binding proteins of the calreticulin protein family (CALR, CANX) (Davidi *et al*, 2020) and gelsolin (GSN). When we assessed the overlap of our data with proteins previously identified in LBs from PD cases with dopaminergic neuronal loss (NL) (Petyuk *et al*, 2021), we found that 17 out of 156 overlapping proteins detected overall showed structural changes upon fibril spike-in, with a significant enrichment (*p*-value < 0.01, Fisher’s exact test) (Figure S 1d). Within this set of 17 proteins, we detected SEC31A (SEC31 Homolog A, COPII coat complex component), which is involved in ER to Golgi transport and macroautophagy (Antoniou *et al*, 2022) and was shown to exhibit altered protein levels in neurons. In particular, we also identified peptidyl-prolyl cis-trans isomerase FKBP1A, which was observed to promote the aggregation of aSyn and cause abnormal, nonlinear, hydrophobic aggregation of aSyn (Caminati & Procacci, 2020).

Functional enrichment analysis (*q*-value < 0.01, Benjamini-Hochberg FDR, minimum hypergeometric test, SimRel functional similarity, size = 0.7) on the set of putative fibril interactors showed enrichment in purine ribonucleoside triphosphate binding, including ATP- and GTP-binding proteins, proteins with catalytic activity, and cytoskeletal protein binding (molecular functions), in extracellular membrane-bounded organelles, vesicles, microtubules, or cytoskeleton (cellular components) and in the regulation of cellular component organization and macromolecule localization (biological processes). Several of these terms correspond to processes and pathways known to be modulated in neurodegenerative diseases, including PD.

Next, we asked whether the proteins we identified as structurally altered upon spike-in of monomeric or fibrillar aSyn could be physically or functionally linked to disease-associated genes that modulate the probability of developing PD, based on genome-wide association studies (GWAS) and also including rare variants implicated in disease. For this, we used a comprehensive interaction network consisting of physical or functional interaction data and performed network-based expansion (the personalized PageRank algorithm) followed by walktrap clustering for monomer- and fibril-interacting proteins and known PD-associated genes, enabling the identification of gene modules and shared biological processes that overlap with PD-associated genes, as described previously (Barrio-Hernandez *et al*, 2021). Within our set of structurally altered proteins, we found four significant modules (Figure 4g) enriched for genes involved in biological processes that are associated with common or rare variants of PD-associated genes (*q*-value < 0.05, Benjamini-Hochberg FDR, Fisher’s exact test, one-sided) (Figure 4h), specifically protein polyubiquitination (module 1), ATP metabolism (module 2), RNA splicing/mRNA processing (module 3), and receptor-mediated endocytosis (module 4). We only identified modules corresponding to protein polyubiquitination and ATP metabolism upon addition of fibrillar but not monomeric aSyn, indicating that these processes may be impaired in PD via interactions with aSyn amyloid fibrils. On the contrary, endocytotic processes, such as synaptic vesicle recycling and clathrin-dependent endocytosis, were specifically enriched for aSyn monomer. In summary, these results show that our method can identify aSyn conformation-specific structurally altered proteins that are linked to PD.

Due to the pathological relevance of aSyn amyloid fibrils, we further probed putative fibril-specific interactions using an orthogonal approach. We spiked either monomeric or fibrillar aSyn, or a PBS control, into SNCA-KO iPSC extracts and used quantitative MS to identify proteins that co-precipitate with the insoluble aSyn fibrils upon ultracentrifugation. As expected, aSyn monomer was predominantly recovered in the soluble fraction and Syn fibrils in the pellet. We identified 574 proteins (|FC| > 1.5, moderated t-test, *q*-value < 0.05) that were either enriched in the pellet exclusively upon fibril spike-in or depleted from the supernatant upon spike-in of aSyn monomer or PBS control. These proteins may be direct binders of aSyn fibrils and were indeed enriched in the set of proteins with fibril-dependent structural changes identified by LiP–MS (*p*-value < 0.01, Fisher’s exact test) (Figure S 4e). The overlap between putative aSyn binding proteins in the two datasets is relatively small (53 proteins; approximately 24 % of the identified LiP hits), as expected due to the experimental differences between LiP–MS and ultracentrifugation. With LiP–MS, we probe protein structural changes that can result from direct interactions of aSyn fibrils with other proteins in the lysate irrespective of their binding affinity, or from other indirect structural effects. In contrast, low-affinity interactions may be disrupted during ultracentrifugation. Consistent with this, putative aSyn fibril interactors detected by both techniques generally have a steeper dose-response curve compared to those identified only by LiP–MS; this may indicate higher relative binding affinity (but see Discussion). Candidate binders identified by both assays are likely high-confidence interactors and are provided (Supplementary Data 4).

Taken together, our LiP–MS approach identified several known as well as novel putative interactors of aSyn. Although some of these have previously been shown to exclusively interact with aSyn monomer or fibrils, in most previous studies it is not clear if proteins are interactors of the monomeric or the aggregated protein and aSyn conformation-dependent interactions were typically probed only for a few proteins (van Diggelen *et al*, 2020; Betzer *et al*, 2015; Leitão *et al*, 2021). In comparison, our results demonstrate that we can systematically compare which interactions occur with either of the structural states of aSyn *in situ* and, most importantly, directly in complex cellular extracts without prior labeling and purification. Our study thus provides a dataset of putative interactors for the monomeric and fibrillar states of aSyn, including interaction interfaces, that will be a rich resource for future follow-up studies.

## Discussion

Interactomics studies remain very challenging for proteins that adopt multiple structural states within the cell. Here, we present a LiP–MS-based structural proteomics approach that enables the identification of structure-specific putative interactomes of any protein that can be purified in a defined state or controllably switched between states. We validated the ability of LiP–MS to map PPIs between purified proteins as well as between proteins in complex biological matrices, and applied the approach to identify differential interactomes for the monomeric and fibrillar states of the PD-associated protein aSyn. We have shown that LiP–MS identifies altered proteolytic patterns upon protein–protein binding and that its peptide-level resolution enables the identification of PPI interfaces *in situ*. Knowing interaction interfaces is useful for structural characterization of protein complexes, the introduction of mutations to disrupt interactions, and the potential development of drugs that target specific PPIs of interest.

Our structural proteomics approach should enable the derivation of quantitative binding parameters, such as relative affinities, for PPIs directly in complex cell lysates. This capability has been previously demonstrated for protein–small molecule interactions using a closely related method, LiPQuant (Piazza *et al*, 2020). This will then enable the ranking of PPIs based on *in situ* relative affinity, which can be used to prioritize targets for further investigation. However, the validity of this approach will require further studies.

Our demonstration that LiP–MS detects interactions between site-specific antibodies and conformers of RSVF has implications in the field of antibody development beyond validation of our method for the detection of PPIs. In experiments with purified proteins, our method pinpointed the exact locations of the three well-characterized antigenic sites on prefusion and postfusion RSVF conformers. We detected no interactions between antigenic site Ø-specific antibodies and postRSVF. This was expected since this antigenic site is known to be solvent-inaccessible in the postfusion conformation. This ability to accurately detect site- and conformation-specific interactions between antibodies and target proteins should be useful for the identification of structure-specific antigenic sites as well as the characterization of novel antibodies, in particular when high-resolution structures are difficult to obtain. Importantly, the dose-response experiment in which a site-specific antibody was added to whole cell extracts was able to identify the specific target interaction, interaction interface, and relative protein binding parameters as previously demonstrated for small molecules (Piazza *et al*, 2020). This type of experiment is expected to become a tool for screening for off-target binders in both basic and pharmaceutical research.

Furthermore, we demonstrated that protein interaction measurements can be performed on crude membrane suspensions using LiP in combination with label-free quantitative MS. Importantly, we showed that protein sequence coverage of membrane proteins is significantly improved when utilizing crude membrane suspensions compared to standard cellular extracts, from which membrane proteins are largely removed. Our approach enabled studying PPIs that are difficult to resolve by other structural proteomics methods such as cryo-EM or X-ray crystallography. As such, our method can be used to better understand the participation of unstructured and flexible regions in PPIs, thus gaining deeper insights, for example, into the regulation and mechanism of action of ACs or other proteins with such domains. We were able to detect known interaction interfaces between AC8 and CaM including their relative binding parameters *in situ*, and our system-wide analysis identified other putative CaM-binding proteins that contain canonical CaM-binding motifs. These novel putative interactors should be characterized in future studies. Our proof-of-principle study demonstrates that LiP in combination with label-free quantitative MS will be very valuable for the analysis of IMPs as these targets are typically difficult to purify and immunoprecipitate.

Using the LiP–MS structural readout, we have probed the interactome of monomeric and fibrillar forms of aSyn, which undergoes protein misfolding and aggregation events and localizes in brain deposits, called LBs, in individuals with the disease. Previous studies have suggested that the interactome of aSyn can be conformation-dependent; however, conformation-specific interactions were demonstrated for only a small subset of proteins (Betzer *et al*, 2015; Lassen *et al*, 2016; Leitão *et al*, 2021; van Diggelen *et al*, 2020). Our work extends these prior studies to a proteome-wide scale, shows that aSyn monomers and fibrils likely interact with different sets of cellular proteins, and provides the rich resource of an extensive putative interactome of these aSyn conformations *in situ*.

In the course of our study, we identified a number of previously described interactors of aSyn, some of which are known to specifically interact with the monomeric form (CALM1) or fibrillar form (e.g., HSP90AA1, HSP90AB1, HSPA1A, HSPA8, and RAB3A) of the protein. However, other known interactors of aSyn (e.g., PINK1, PARK2, LRRK2) were not detectable in our analysis, primarily due to their low expression levels in neurons, which is in accordance with previous literature (Lee *et al*, 2020). This reflects a general caveat of our approach, which relies on MS detection and sufficient sequence coverage of proteins in order to be able to investigate them as potential interactors.

We observed many more structurally altered proteins upon addition of aSyn fibrils than upon addition of monomer, consistent with the known lower binding preference of the aSyn monomer (Leitão *et al*, 2021; Betzer *et al*, 2015; van Diggelen *et al*, 2020). Notably, we have identified novel aSyn monomer-specific structural alterations on proteins involved in RNA binding and protein binding. We further identified putative interactors localized in extracellular organelles, cell junctions, vesicles, as well as mitochondria. In contrast, we detected a set of fibril-specific putative interactors, including ATP- and GTP-binding proteins, proteins with catalytic activity, and cytoskeletal protein binding. These proteins localize in extracellular vesicles, vesicles, or microtubules, some of which were previously unknown interactors and would require follow-up to confirm direct aSyn conformation-specific interaction. Interestingly, a subgroup of our potential aSyn fibril interactors are components of LBs or implicated in their formation (Xia *et al*, 2008; Petyuk *et al*, 2021; Mahul-Mellier *et al*, 2020). Although LB formation is a complex process, it is possible that this set of proteins interacts with fibrillar aSyn also in LB *in vivo*.

The network-based propagation and clustering analysis demonstrate disease-relevant links to common or rare variants of PD, such as involvement of protein polyubiquitination, ATP metabolism, RNA splicing/mRNA processing, and receptor-mediated endocytosis. One interesting aspect that emerged from the analysis of PD-associated genes was that fibril specificity was observed for intracellular energy metabolism and protein polyubiquitination. In this regard, defective mitochondrial functions that lead to increased oxidative stress have been demonstrated to play a central role in PD pathogenesis (Hattori & Mizuno, 2015). In particular, deficiencies of mitochondrial respiratory chain complex I may lead to the degeneration of neurons in PD by reducing the synthesis of ATP. In our analysis, we revealed interactions of aSyn fibrils with three proteins (NDUFB5, NDUFS1, and NDUFV2) that are part of the mitochondrial respiratory chain complex I. Furthermore, two proteins (ATP5PB and UQCRFS1) that showed structural alterations are part of the inner mitochondrial membrane protein complex. It is suggestive that, contrary to physiological, monomeric aSyn, pathogenic aSyn fibrils preferentially bind to mitochondria and cause mitochondrial respiration defects, as proposed by previous studies (Wang *et al*, 2019).

Growing evidence strongly implicates a direct role of the ubiquitin-proteasome system (UPS) in the pathogenesis of PD (Lim & Tan, 2007). The UPS is a type of intracellular protein degradation machinery and its disruption in the presence of aSyn fibrils could lead to dysfunction of associated protein quality control mechanisms. We identified associations of three structurally altered proteins (CAND1, RCHY1, and UBE2O) with a common variant of the *PRKN* gene linked to PD. This gene encodes for the Parkin protein that functions as an E3 ubiquitin ligase. Interestingly, aggregated aSyn has been shown to selectively interact with the 19S cap and concomitantly inhibit the function of the 26S proteasome (Snyder *et al*, 2003).

In regard to endocytotic processes, we observed associations with genes involved in synaptic vesicle recycling and clathrin-dependent endocytosis. Through our analysis, we identified structurally altered proteins, specifically SYNJ1 and PACS1, that were found in the module of genes associated with common variants linked to PD. Previous studies provide evidence that an excess of aSyn monomer impairs clathrin-mediated synaptic vesicle endocytosis, as indicated by a loss of synaptic vesicles (Medeiros *et al*, 2018).

In our study, we utilized recombinantly expressed aSyn and amyloid fibrils generated *in vitro*. While previous studies have employed *in vitro* generated aSyn amyloid fibrils to study aSyn aggregation (Narkiewicz *et al*, 2014; Viennet *et al*, 2018; Afitska *et al*, 2020), it is important to note that the structures of amyloid fibrils produced in the test tube may differ from those found *in vivo* fibrils in the brains of individuals with PD. Recent studies have generated brain-derived aSyn fibrils by amplification from brain extracts (Strohäker *et al*, 2019). These could be used in our approach to studying aSyn interactors in PD, but also in other synucleinopathies, such as DLB or MSA. It should be noted that aSyn fibrils from different synucleinopathies are thought to adopt distinct conformations, known as amyloid strains. Our approach could thus enable the analysis of strain-specific interactomes for different strains of aSyn fibrils linked to various clinical phenotypes. Similarly, although we chose to use lysates of human SNCA-KO iPSC-derived cortical neurons in our experiments in order to reduce the potential background from endogenous aSyn, our approach could be applied to extracts of any other cellular or organismal model of PD.

A key advantage of our approach, as we have shown in this study, is that PPIs can be detected by adding a purified protein, in different conformations if desired, directly into complex extracts without the need for prior labeling. However, a few limitations should be noted. Cell lysis can lead to artifacts due to disruption of subcellular organization. Further, the method relies on the purity of proteins introduced into cellular extracts as contaminants could cause structural changes in the extract through direct binding or other indirect effects. In addition, although we have demonstrated that our approach identifies PPI interfaces directly, it may also detect conformational changes in other parts of the protein that occur due to protein binding or indirect effects triggered by binding to the target protein. Therefore, putative interactors must be confirmed using orthogonal methods.

Collectively, our work demonstrates that our LiP–MS approach successfully identifies PPIs *in situ* and detects interaction interfaces of different classes of proteins, including antibodies, membrane proteins, structured proteins, proteins with unstructured, intrinsically disordered regions, and disease-associated, amyloidogenic proteins, which remain difficult to study in classical interactomics experiments. LiP–MS further allows for the profiling of differential interactomes of different structural states of proteins *in situ*. As we demonstrate for aSyn, our method can be applied generally to study the interactomes of disease-relevant proteins that undergo structural changes, and could thus help identify novel targets in drug discovery.

## Supplementary Figures

**Figure S 1:**
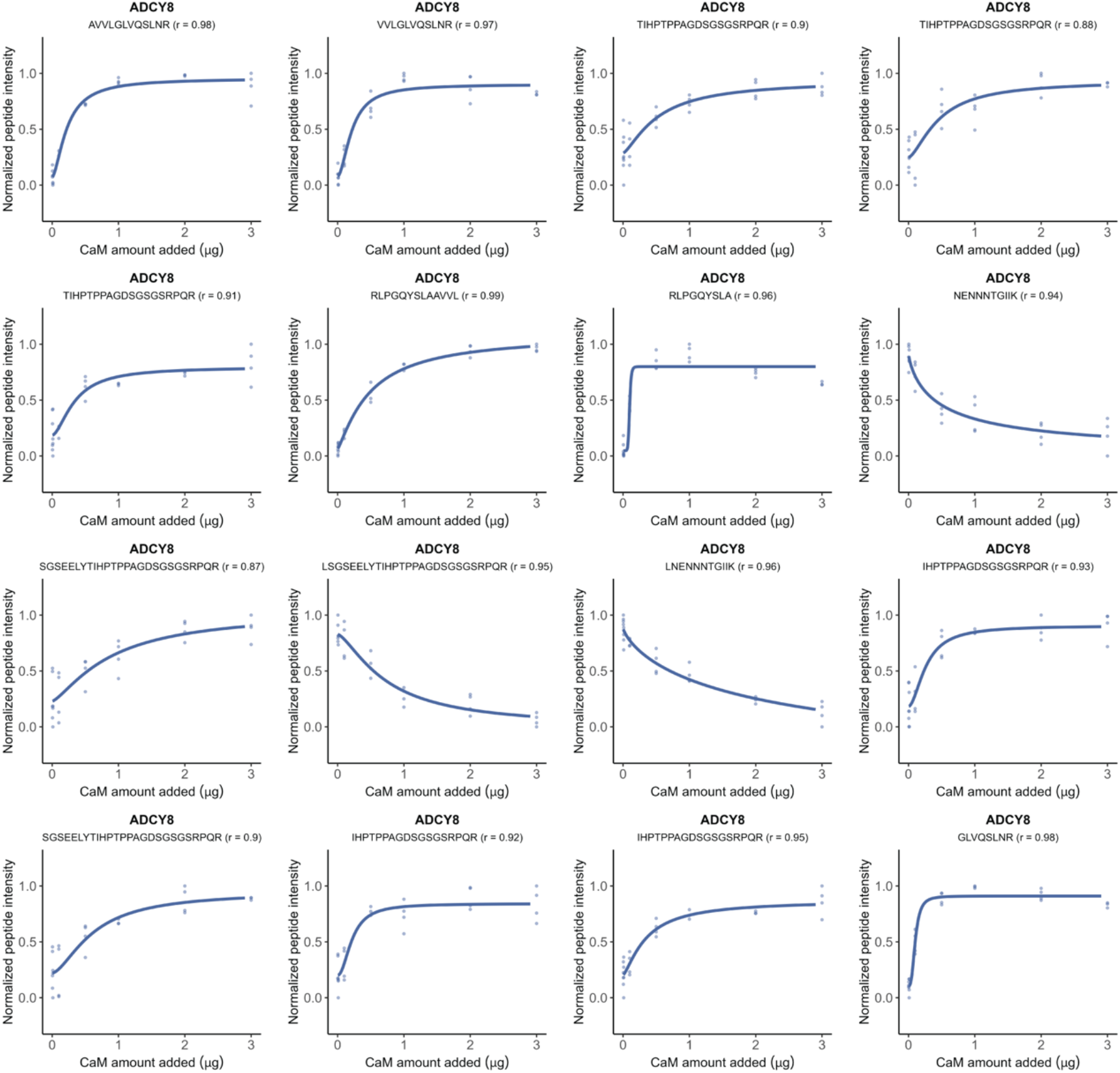
Dose-response curves of sixteen LiP peptides originating from AC8 show relative peptide intensities proportional to the amount of CaM spiked into crude membranes. Pearson’s coefficient (*r*) to a sigmoidal trend of the peptide-intensity response profile is indicated.

**Figure S 2:**
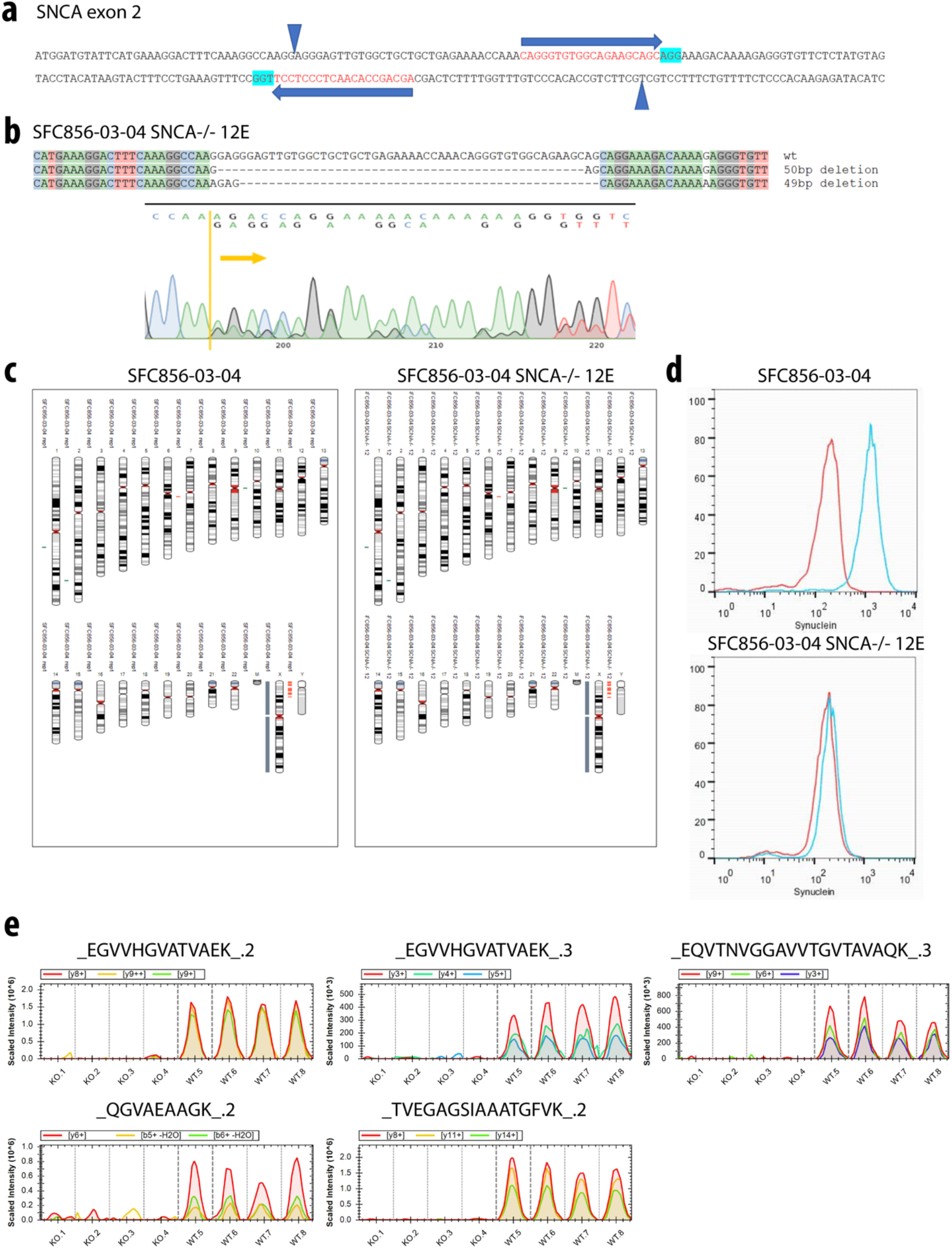
CRISPR/Cas9-mediated KO of SNCA from human iPSC and characterization of previously unpublished derived KO clone SFC856-03-04 SNCA−/− 12E. (a) The editing strategy for creating SNCA-KO from the healthy donor iPSC line SFC856-03-04, shows the exon 2 sequence section with guide RNA positions (blue arrows), PAM (highlight in turquoise), and expected cut positions (blue triangles). (b) The deconvoluted sequence of both alleles of clone 12E shows out-of-frame repair that will lead to premature translational termination, with the original sequence trace in the lower panel. (c) Karyograms produced from the SNP array show no gross abnormalities in clone SFC856-03-04 SNCA−/− 12E versus the parental iPSC line SFC856-03-04. Red indicates loss or single copy, green indicates gain of copy, and grey indicates loss of heterozygosity on autosomes or two copies of the X chromosome (*i.e*., female line). (d) Flow cytometry of macrophages differentiated from iPSC with and without SNCA KO (left and right panels, respectively), stained for aSyn (MJFR1 antibody – blue line, isotype control – red line). (e) MS2 extracted ion chromatographs for five peptide precursors of aSyn quantified in SNCA-KO (n = 4) and healthy control (WT; n = 4) iPSC lines. The color coding of the fragments is indicated.

**Figure S 3:**
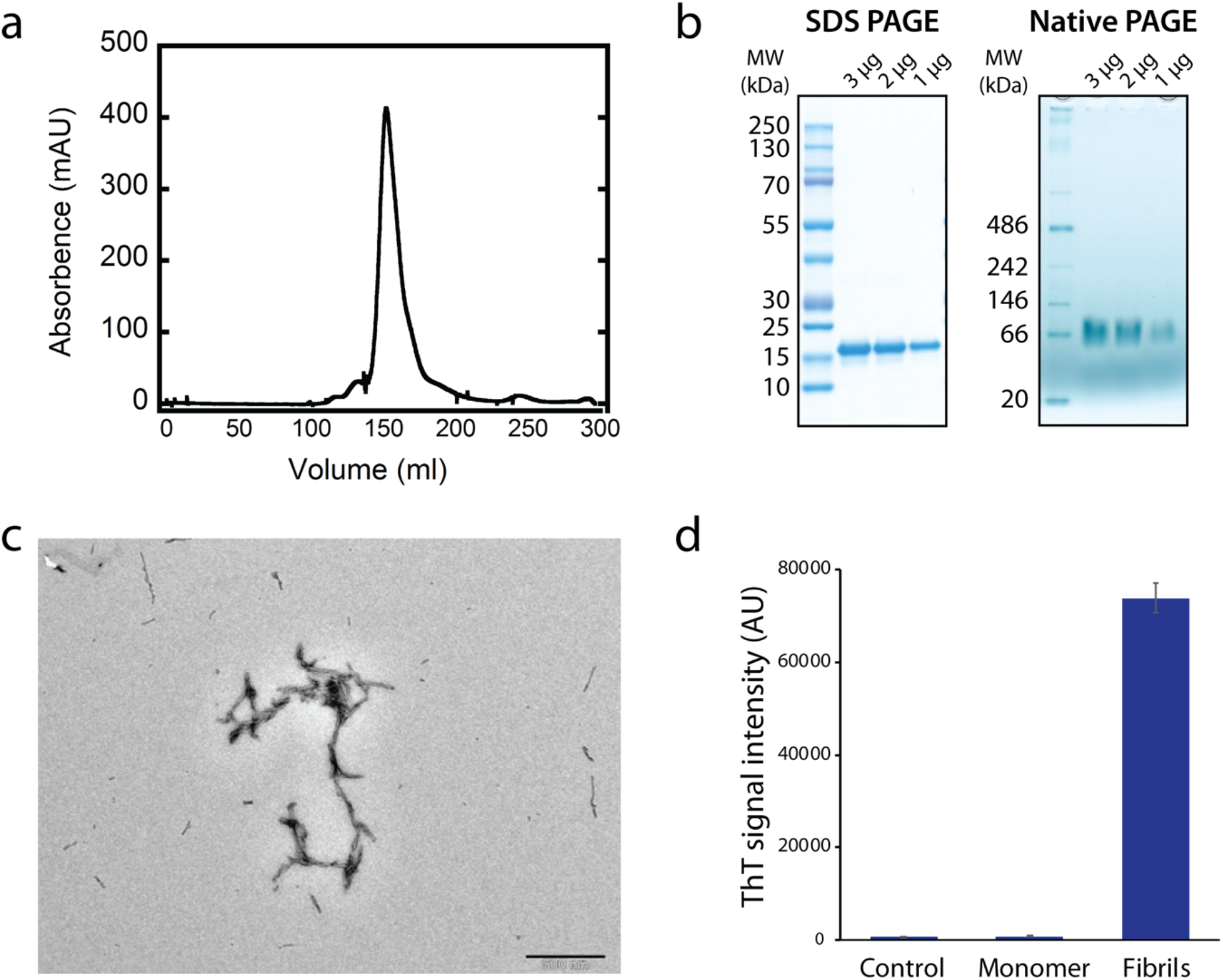
Quality control experiments for aSyn monomer and amyloid fibrils. (a) SEC analysis of purified aSyn monomer. (b) SDS PAGE and Native PAGE of monomeric aSyn. (c) TEM image of aSyn amyloid fibrils. (d) ThT intensity of PBS control, aSyn monomer, and aSyn amyloid fibrils.

**Figure S 4:**
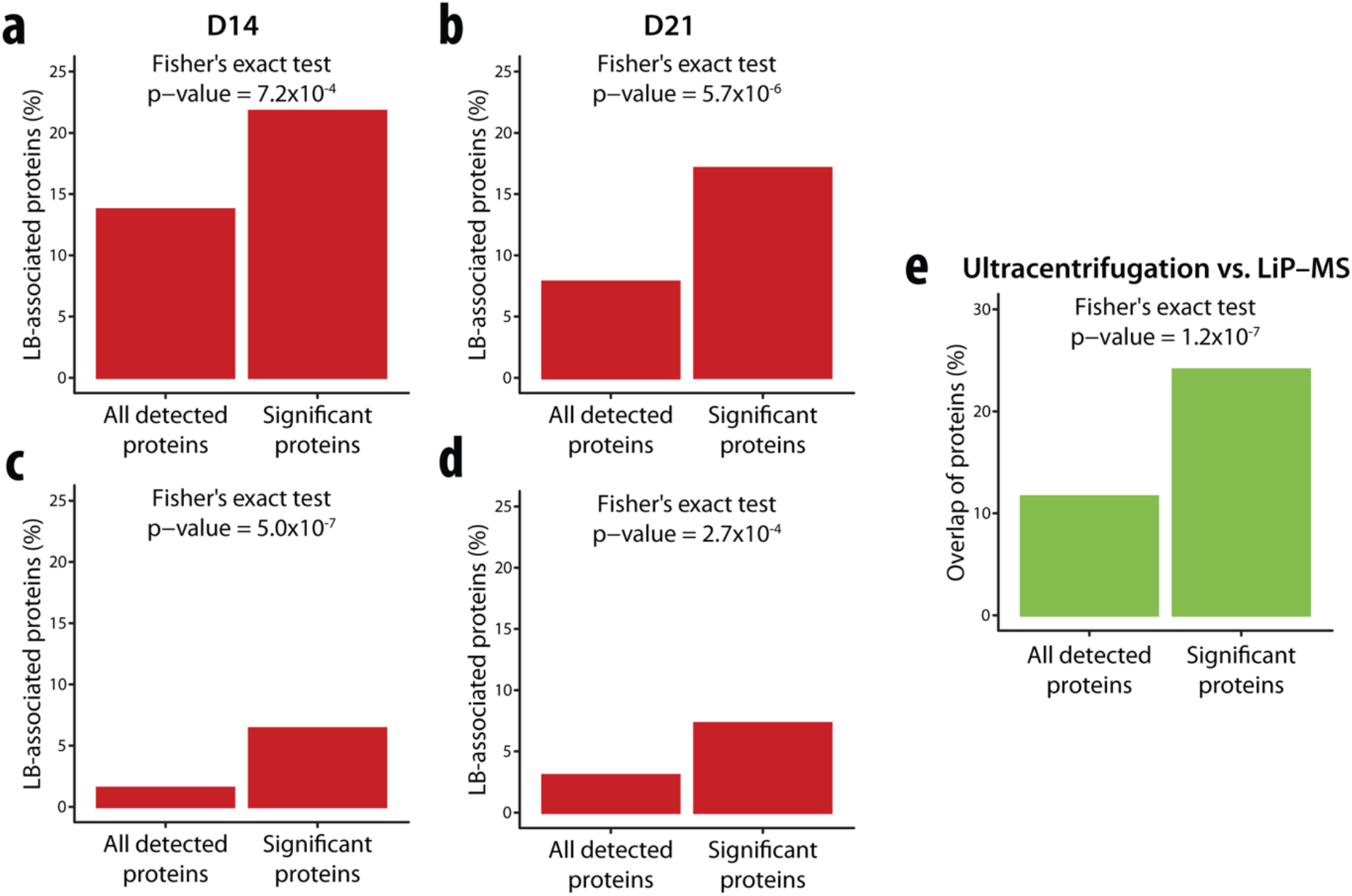
Identification of Lewy body (LB)-associated and precipitated proteins. (a-d) and fibril-binding proteins by ultracentrifugation (e). (a, b) Enrichment analyses of putative LiP–MS-identified fibril-binding proteins for proteins previously identified as LB components in a neuronal seeding model at two-time points (a, day 14; b, day 21) (Mahul-Mellier *et al*, 2020). The plots show the fraction of previously identified LB components in structurally altered proteins (right bar) versus all detected proteins (left bar) upon spike-in of aSyn fibril into an iPSC-derived cortical neuron lysate. *p*-values assessing enrichment (Fisher’s exact test) are shown. (c, d) Enrichment analysis as in a, b for proteins detected in patient-derived LBs (Xia *et al*, 2008) (c) and for proteins identified in patient-derived LBs with neuronal loss (Petyuk *et al*, 2021). (e) Enrichment analysis of putative LiP– MS-identified fibril-binding proteins for ultracentrifugation-identified putative fibril binders. The plots show the fraction of ultracentrifugation-identified fibril binders in structurally altered proteins (right bar) versus all detected proteins (left bar) upon spike-in of aSyn fibril into an iPSC-derived cortical neuron lysate. *p*-values assessing enrichment (Fisher’s exact test) are shown.

## Supplementary Tables

**Supplementary Table 1:** Definition of variable windows for DIA–MS measurements.

| Window | Mass-to-charge ratio ($m/z$) | Charge ($z$) | Isolation window ($m/z$) |
| --- | --- | --- | --- |
| 1 | 358 | 2 | 16 |
| 2 | 373 | 2 | 16 |
| 3 | 388 | 2 | 16 |
| 4 | 403 | 2 | 16 |
| 5 | 418 | 2 | 16 |
| 6 | 433 | 2 | 16 |
| 7 | 448 | 2 | 16 |
| 8 | 463 | 2 | 16 |
| 9 | 478 | 2 | 16 |
| 10 | 493 | 2 | 16 |
| 11 | 508 | 2 | 16 |
| 12 | 523 | 2 | 16 |
| 13 | 538 | 2 | 16 |
| 14 | 553 | 2 | 16 |
| 15 | 568 | 2 | 16 |
| 16 | 583 | 2 | 16 |
| 17 | 598 | 2 | 16 |
| 18 | 613 | 2 | 16 |
| 19 | 628 | 2 | 16 |
| 20 | 643 | 2 | 16 |
| 21 | 659 | 2 | 18 |
| 22 | 676 | 2 | 18 |
| 23 | 693 | 2 | 18 |
| 24 | 710 | 2 | 18 |
| 25 | 727 | 2 | 18 |
| 26 | 744 | 2 | 18 |
| 27 | 761 | 2 | 18 |
| 28 | 778 | 2 | 18 |
| 29 | 795 | 2 | 18 |
| 30 | 813 | 2 | 20 |
| 31 | 832 | 2 | 20 |
| 32 | 851 | 2 | 20 |
| 33 | 870 | 2 | 20 |
| 34 | 889 | 2 | 20 |
| 35 | 908 | 2 | 20 |
| 36 | 929.5 | 2 | 25 |
| 37 | 953.5 | 2 | 25 |
| 38 | 977.5 | 2 | 25 |
| 39 | 1006.5 | 2 | 35 |
| 40 | 1048 | 2 | 50 |
| 41 | 1111 | 2 | 78 |

## Materials and Methods

### Experimental model and subject details

#### HEK293T cells

HEK293T cells were cultured in Dulbecco’s Modified Eagle’s Medium (DMEM) (Thermo Fisher) supplemented with 10% fetal bovine serum (FBS) and 1% penicillin/streptomycin. The HEK293T cells were passaged prior to confluency by detachment with 0.25% trypsin, followed by two consecutive washing steps in LiP buffer (100 mM HEPES pH 7.5, 150 mM KCl, 1 mM MgCl_2_). To store the pellets, the HEK293T cells were transferred to 1.5-mL Eppendorf tubes and centrifuged at 1,000 x g at 4 °C for 5 minutes. LiP buffer was removed and the pellets were snap frozen and stored at –80 °C until further use.

#### HEK293F membranes with AC8 overexpression

The full-length DNA construct of bovine AC8 was cloned into a tetracycline-inducible pACAMV-based vector with a C-terminal HRV 3C cleavage site and YFP-Twin-Strep fusion tag. The plasmid was transfected into HEK293F GnTI- cells and a stable clone expressing AC8 was selected for further protein expression.

#### CRISPR/Cas9-mediated knockout of SNCA in human induced pluripotent stem cells (iPSCs)

The previously published human iPSC line SFC856-03-04 was used for gene editing (Haenseler *et al*, 2017b). The iPSC line was derived from a healthy donor (78y female) and is karyotypically normal as assessed by SNP analysis. SNCA has 6 exons, exon 2 is the first coding exon, and exons 3 and 5 can be alternatively spliced out. Exon 2 was chosen as the target for the introduction of a 49bp deletion (downstream from the translation start site) that would lead to a frameshift and subsequent premature stop codon. The dual guide RNA sequences and double-strand cut strategy is illustrated in Figure S 2a, and utilized Alt-R CRISPR-Cas9 crRNA/trRNA/Hi-Fi Cas9 ribonucleotide-protein complex (IDT) and Neon electroporation (ThermoFisher) to deliver the complex to the iPSC. Clones were screened by PCR and those clones harboring the deletion were sequenced across the repair to confirm out-of-frame repair (Figure S 2b). Clone SFC856-03-04 SNCA−/− 12E was used for the experiments presented here, it was confirmed to still retain the same karyotype as the parent iPSC line by Illumina Ominexpress24 SNP array and pseudokaryogram visualization using Karyostudio (Figure S 2c). The knockout of SNCA at the protein level was confirmed by differentiation to macrophages (Haenseler *et al*, 2017b), which express readily measurable levels of alpha-synuclein by flow cytometry (Figure S 2d); antibody MJFR1 (Abcam 138501) was used to detect aSyn, alongside a matched isotype control.

#### iPSC culture and differentiation to cortical neurons

The healthy control SFC856-03-04 and the edited SFC856-03-04 SNCA−/− 12E knockout iPSC lines were used in the study presented here. iPSCs were cultured in Essential8 medium (Thermo Fisher) on Geltrex-coated tissue culture plates and passaged as small cell clusters using 0.5 mM EDTA (Thermo Fisher). iPSCs were differentiated into cortical neural progenitor cells (NPCs) with a dual SMAD inhibition protocol (Shi *et al*, 2012), with minor modifications as described previously (Haenseler *et al*, 2017a). NPCs were frozen on differentiation day 28 or directly used for further differentiation and experiments. The final plating of cells was on differentiation day 36. To remove proliferating progenitors and astrocytic cells cultures were treated with 2 μM AraC (Sigma Aldrich) for 3 days. Experiments were performed on differentiation day 56. *Respiratory syncytial virus fusion (RSVF) glycoprotein*: The construct encoding the stabilized prefusion RSVF glycoprotein, known as DS2 (Joyce *et al*, 2016), corresponds to the sc9-10 DS-Cav1 A149C Y458C S46G E92D S215P K465Q variant. The codon-optimized sequence for mammalian cells was expressed and cloned into the pHCMV-1 vector flanked with two C-terminal Strep-Tag II and one 8x His tag. Expression and purification were performed as described previously (Sesterhenn *et al*, 2020). The postfusion RSVF glycoprotein was expressed, purified, and provided by Fabian Sesterhenn.

#### RSVF site-specific antibodies

All site-specific monoclonal antibodies against RSVF used in this study were expressed, purified, and provided by Fabian Sesterhenn. Prior to LiP experiments, all antibodies were diluted in PBS buffer, pH 7.0 to a concentration of 1 μg/μL. For LiP titration experiments, a total amount of 0.5, 1.0, 1.5, 2.0, 3.0, and 4.0 μg of motavizumab was spiked into each sample containing 100 μg of total proteins.

#### Human IgG1 kappa antibody

The InVivoMAB human IgG1 isotype control antibody (BioXCell) purified from human myeloma serum was diluted in PBS buffer, pH 7.0 to a concentration of 1 μg/μL and used for subsequent LiP experiments with purified RSVF.

#### Calmodulin (CaM)

Lyophilized CaM from bovine testes (Sigma Aldrich) was solubilized in LiP buffer (100 mM HEPES pH 7.4, 150 mM KCl, 1 mM MgCl_2_). For the CaM titration experiment, CaM was spiked into crude membranes at an amount of 0.01, 0.1, 0.5, 1, 2, and 3 μg per 100 μg of total crude membrane proteins.

#### Alpha-synuclein (aSyn) monomer

N-terminally acetylated, human wild-type aSyn was expressed in *E. coli* cells (strain BL21 Star, DE3) transfected with the pRK172 plasmid with the yeast N-acetyltransferase complex B (NatB) as described previously (Kumari *et al*, 2021). Protein purification was performed as described elsewhere (Campioni *et al*, 2014). Lyophilized aSyn was resuspended in PBS buffer, pH 7.4. Subsequently, spun down with ultracentrifugation at 100,000 x g for 30 min before sample preparation to separate oligomeric species from monomeric aSyn. For LiP experiments, monomeric aSyn was spiked into iPSC SNCA-KO cell extracts at an amount of 0.05, 0.1, 0.5, 1.0, 2.0, and 4.0 μg per 50 μg of total proteins.

#### aSyn amyloid fibrils

For the formation of mature aSyn amyloid fibrils, Eppendorf LoBind microcentrifuge tubes (1.5 mL) containing 0.75 mL of 5 mg/mL monomeric aSyn in PBS buffer, pH 7.4, 150 mM NaCl were incubated at 37 °C on a thermomixer under agitation at 800 rpm for 1–2 weeks. Amyloid fibrils were sonicated to produce shorter fibrillar structures as described elsewhere (Patterson *et al*, 2019). To remove low-molecular species, amyloid fibrils were pelleted by ultracentrifugation at 100,000 x g for 30 min and diluted in fresh PBS buffer, pH 7.4 to a concentration of 1 μg/μL. Subsequently, 0.05, 0.1, 0.5, 1.0, 2.0, and 4.0 μg of amyloid fibrils were added to each sample containing 50 μg of total proteins of the SNCA-KO iPSC extract.

### Thioflavin T (ThT) fluorescence assay

The ThT binding assay was performed to confirm the presence of aSyn amyloid fibrils using a 40 μM ThT solution. Aliquots of aSyn monomer and amyloid fibrils were added to the ThT solution in three replicates and fluorescence emission was measured at 25 °C on a CLARIOstar Plus plate reader (BMG Labtech) with an excitation wavelength of 440 nm. Fluorescence emission was recorded at a wavelength of 484 nm.

### Transmission electron microscopy (TEM)

Samples of amyloid fibrils were examined by TEM with negative staining. A droplet of the sample was placed on carbon film-coated copper grids, dried, and negatively stained with a droplet of 1% (w/v) uranyl acetate. The TEM images of the amyloid fibrils were imaged using a Hitachi HT7700.

### Blue Native Polyacrylamide Gel Electrophoresis (BN-PAGE)

BN-PAGE was performed using NativePAGE™ Sample Prep Kit and precasted NativePAGE™ 4 to 16%, Bis-Tris gels (1.0 mM, Mini Protein Gel, 10-well). Samples containing 1, 2, and 3 μg of monomeric aSyn were diluted with NativePAGE™ 4X Sample Buffer. Samples and the NativeMark™ Unstained Protein Standard were loaded into wells filled with 1X NativePAGE™ Dark Blue Cathode buffer, containing Coomassie G-250. Gels were run at 150 V constant in NativePAGE™ Dark Blue Cathode buffer at the Cathode and NativePAGE™ Anode buffer at the Anode for 30 minutes. NativePAGE™ Dark Blue Cathode buffer was exchanged with NativePAGE™ Light Blue Cathode buffer. The gel was run until completion at 150 V constant. Gels were fixed in fix solution (40 % methanol, 10 % acetic acid) and microwaved for 45 seconds, followed by shaking on an orbital shaker for 15 minutes. The gels were then destained in destaining solution (8 % acetic acid) and microwaved for 45 seconds, followed by incubation on the orbital shaker for 15 minutes. This procedure was repeated multiple times until the gel was completely destained.

### SDS-PAGE

SDS-PAGE was performed using precasted 4–12% NuPAGE™ Bis-Tris gels in NuPAGE™ MES SDS Running Buffer. Laemmli buffer (5x) was added to the samples containing 1 μg, 2 μg and 3 μg monomeric aSyn. As a marker, we used PageRuler Plus Prestained protein ladder. The gel was run at 80 V constant for 15 minutes, followed by 150 V constant until completion. The gels were stained using PageBlue™ Protein Staining Solution, and destaining was achieved by shaking on an orbital shaker in double-deionized water.

### Preparation of cell extracts for MS analysis

#### HEK293T and iPSC-derived cortical neurons

All steps throughout sample preparation were performed on ice. Cell pellets were resuspended in 400 μL LiP buffer (100 mM HEPES pH 7.4, 150 mM KCl, 1 mM MgCl_2_) and lysed using a pellet pestle (Argos Technologies) in ten cycles of 10 s of homogenization and 1-min pause at 4 °C. The lysate was cleared by centrifugation (1,000 x g at 4 °C) for 15 min. The supernatant was transferred to a new Eppendorf tube and the remaining pellet was further resuspended in 200 μL LiP buffer. The lysis step was repeated as described and supernatants were combined. The total lysate protein concentration was determined with a Pierce BCA Protein Assay Kit (cat #23225) according to the manufacturer’s instructions.

### Preparation of crude membranes for MS analysis

All steps throughout crude membrane preparation were performed on ice. Pellets of 2L HEK293F GnTI- cells overexpressing bovine AC8 were resuspended in 50 mL LiP buffer (100 mM HEPES pH 7.4, 150 mM KCl, 1 mM MgCl_2_) supplemented with one tablet of Roche cOmplete EDTA free inhibitor cocktail and 0.01 mg/mL DNAse. Cells were lysed using a dounce homogenizer with 20 strokes and centrifuged at 1,000 x g at 4 °C for 10 min. The supernatant was split, transferred to two ultracentrifuge tubes containing 25 mL LiP buffer containing protease inhibitors and DNAse, and spun down (Ti45 rotor, 35,000 rpm at 4 °C) for 40 min. The remaining pellets were resuspended in 15 mL LiP buffer and further homogenized using a dounce homogenizer with 20 strokes. The total protein concentration of crude membranes was determined with a Pierce BCA Protein Assay Kit (cat #23225) according to the manufacturer’s instructions. Crude membranes were stored at –80 °C prior to further use.

### Limited proteolysis in native conditions

Purified RSVF proteins were incubated with site-specific and human isotype control antibodies at a molar ratio of 1:1 (protein:antibody) for 10 min at 25 °C and subjected to limited proteolysis. Proteinase K (PK) from *Tritirachium album* (Sigma Aldrich) was added simultaneously to all four independent replicates of protein samples per condition (n = 4 for all experiments) with the aid of a multichannel pipette, at an enzyme-to-substrate ratio of 1:100 (w/w) and incubated at 25 °C for 5 min. Proteolytic reactions were stopped by heating samples for 5 min at 99 °C in a heat block. Subsequently, samples were transferred to Eppendorf tubes containing an equal volume of 10% sodium deoxycholate (Sigma Aldrich).

For validation experiments with postRSVF and motavizumab in HEK293 cellular extracts, three independent replicate samples (n = 3) per condition containing 100 μg of the HEK293T cellular extract, supplemented with 1 μg of postfusion RSVF, were exposed to a 5-dosage concentration series of motavizumab (0.5, 1.0, 1.5, 2.0, 3.0, and 4.0 μg). Upon 10-min incubation at 25 °C, the samples were then subjected to limited proteolysis as described above. Additionally, untreated control and one condition of treated samples in this experiment were subjected to trypsin digestion only to control for potential protein abundance changes.

Crude membranes were aliquoted in equivalent volumes for each of four independent replicates (n = 4) containing 100 μg of proteins and incubated with calmodulin at given concentrations. Similarly, aSyn in its monomeric and fibrillar, aggregated state was added at given concentrations to each of four independent replicates (n = 4) for each condition containing 50 μg of iPSC extract. Upon 10-min incubation at 25 °C, the samples were then subjected to limited proteolysis as described above. Additionally, untreated and one condition of treated samples in each experiment were subjected to trypsin digestion only to control protein abundance changes.

### Trypsin digestion in denaturing conditions

Samples from all experiments were reduced with 5 mM tris(2-carboxyethyl)phosphine hydrochloride for 45 min at 37 °C. Alkylation was carried out in 40 mM iodoacetamide followed by incubation at RT in the dark for 30 min. Thereafter, samples were diluted in four volumes of 100 mM ammonium bicarbonate and digested with lysyl endopeptidase and trypsin (both at an enzyme-to-substrate ratio of 1:100) at 37 °C for 16 hours. Digests were acidified by addition of formic acid to a final concentration of 2% and sodium deoxycholate precipitate was removed by filtration using a centrifugation filter at 1,000g for 5 min. Peptides were desalted using a 96-well C18 MACROspin plate with 10-100 μg capacity according to the manufacturer’s instructions. After drying, samples were resuspended in 3% acetonitrile (ACN) and 0.1% formic acid. The iRT kit (Biognosys AG, Schlieren, Switzerland) was added to all proteome samples as instructed by the manufacturer.

### Ultracentrifugation assay

The ultracentrifugation assay was employed to separate proteins interacting with amyloid fibrils of aSyn. SNCA-KO iPSC-derived cortical neuron extracts containing 50 μg of protein were incubated in three independent replicates (n = 3) with 2 μg of aSyn monomer and amyloid fibrils for 10 min at 25 °C and centrifuged at 100,000 x g for 30 min at 4 °C. The supernatant was transferred into a new 1.5-mL Eppendorf tube and the pellet was washed three times with 200 μL of LiP buffer (100 mM HEPES pH 7.4, 150 mM KCl, 1 mM MgCl_2_). The pellet was resuspended in LiP buffer by vortexing for 5 min at RT. The protein concentration of the supernatant and the pellet was determined as described above. The samples were further processed with trypsin digestion in denaturing conditions as described above.

### Mass spectrometry data acquisition

Peptide digests of purified RSVF and antibodies were analyzed in DIA mode on a Thermo Scientific Lumos mass spectrometer (Thermo Fisher) equipped with a nanoelectrospray ion source and coupled to an Easy-nLC 1200 system (Thermo Fisher). Peptides were separated on a 25 cm x 0.75 μm i.d. analytical column (Thermo Fisher) packed with 1.9 μm C18 beads using a linear gradient from 5% to 30% buffer B (95% acetonitrile in 0.1% formic acid) over 30 min and a flow rate of 300 nL/min under ambient conditions. Full MS1 scans were acquired between 350 and 1400 m/z at a resolution of 120,000. The automatic gain control (AGC) target of 8×10^5^ and a maximum injection time of 100 ms were used. Forty-one variable-width windows (Supplementary Table 1) were utilized to measure fragmented precursor ions. DIA-MS2 spectra were acquired at a resolution of 30,000 and an AGC target of 2×10^5^ and an injection time of 54 ms. The normalized collision energy was set to 30.

For RSVF, aSyn, and ultracentrifugation assay proteome samples, peptide digests were analyzed in DDA and DIA modes on a Thermo Scientific Eclipse mass spectrometer (Thermo Fisher) equipped with a nanoelectrospray ion source and coupled to an Easy-nLC 1200 system (Thermo Fisher). Peptides were loaded onto a 40 cm × 0.75 μm i.d. analytical column packed in-house with 1.9 μm C18 beads (Dr. Maisch Reprosil-Pur 120) and separated by a 120 min linear gradient at a flow rate of 300 nL/min with increasing buffer B (95% acetonitrile in 0.1% formic acid) from 3% to 30%. For DDA, a full MS1 scan was acquired over a mass range of 350–1400 m/z at a resolution of 120,000 with an AGC target of 200% and an injection time of 100 ms. DDA-MS2 spectra were acquired at a resolution of 30,000 with an AGC target of 200% and an injection time of 54 ms. To maximize parallelization, a duty cycle time was 3 s. For DIA, a full MS1 scan was acquired between 350 and 1100 m/z at a resolution of 120,000 with an AGC target of 200% and an injection time of 100 ms. Forty-one variable-width windows (Supplementary Table 1) were used to measure fragmented precursor ions. DIA-MS2 spectra were acquired at a resolution of 30,000 with an AGC target of 400% and an injection time of 54 ms. The normalized collision energy was set to 30.

For AC8-CaM proteome samples, 1 μg peptide digests were loaded onto a 40 cm × 0.75 μm i.d. column packed in-house with 1.9 μm C18 beads (Dr. Maisch Reprosil-Pur 120) and separated by a 120 min linear gradient at a flow rate of 300 nL/min with increasing buffer B (95% acetonitrile in 0.1% formic acid) from 3% to 30%. All DIA and DDA runs were acquired on a Thermo Scientific Exploris 480 mass spectrometer (Thermo Fisher). For DDA, a full MS1 scan was acquired between 350 and 1100 m/z at a resolution of 120,000 with an AGC target of 200% and an injection time of 100 ms. DDA-MS2 spectra were acquired at a resolution of 30,000 with an AGC target of 200% and an injection time of 54 ms. To maximize parallelization, a duty cycle time was 3 s. For DIA, a full MS1 scan was acquired between 350 and 1100 m/z at a resolution of 120,000 with an AGC target of 100% and an injection time of 100 ms. Forty-one variable-width windows (Supplementary Table 1) were used to measure fragmented precursor ions. DIA-MS2 spectra were acquired at a resolution of 30,000 and an AGC target of 2000%. The first mass was fixed at 200 m/z and the normalized collision energy was set to 28.

### Mass spectrometry data analysis

Prior to DIA spectra processing, all spectral libraries were generated using the library generation functionality of Spectronaut 15 (Biognosys AG, Schlieren, Switzerland) (Bruderer *et al*, 2015) using the default settings with minor adaptations. In brief, the DIA and/or DDA files were searched against the human UniProt FASTA database (updated 2020-03-20), the MaxQuant contaminants fasta database (245 entries), and the Biognosys’ iRT peptides FASTA database. Raw files for RSVF and AC8-CaM datasets were additionally searched against the prefusion or postfusion RSVF FASTA databases and the AC8 (including tags) FASTA database (uploaded to the public repository), respectively. For LiP–MS datasets, digestion enzyme specificity was set to Trypsin/P and semi-specific. For trypsin-only treated controls, a digestion enzyme was Trypsin/P with specific cleavage rules. The minimum allowed peptide length was set to 5 amino acids with a maximum of two missed cleavages per peptide. Carbamidomethylation of cysteine was considered a fixed modification, and acetylation (protein N-terminus) and oxidation of methionine as variable modifications. DIA spectra were further processed with Spectronaut using the default settings with a few modifications. In short, dynamic retention time extraction was applied with a correction factor of 1. The identification of peptides and proteins was controlled by the false discovery rate (FDR) of 1%. The machine learning algorithm and Q-value calculations were run across the entire experiment. Peptide quantification was carried out on the modified peptide sequence level using precursor ions. Protein quantification included only proteotypic peptides and global median normalization was applied.

### Interpretation of antibody-target protein interactions with purified proteins

Peptide reports provided by Spectronaut were processed using an in-house R script in R Statistical Software (version 4.2.0; R Core Team 2021). Raw abundances of proteotypic RSVF peptides were normalized using variance stabilizing normalization with the *vsn* package (Huber *et al*, 2002) (version 3.64.0). The normalized log_2_-transformed peptide abundances from treated samples with site-specific antibodies against RSVF were compared to control, *i.e*., anti-Human IgG1 kappa antibody with RSVF. The log_2_ FC and the statistical significance (represented by *p*-values adjusted by multiple testing using the Benjamin-Hochberg method) were computed using an empirical Bayes moderated t-test provided by the *limma* package (Ritchie *et al*, 2015) (version 3.52.4). Peptides with at least three measured peptide abundances for each condition, fulfilling the defined criteria (|log_2_ FC| > 1, *q*-value < 0.01), were considered significant. Significant peptides were mapped onto 3D structures of prefusion (PDB: 4JHW) (McLellan *et al*, 2013) and postfusion (PDB: 3RRR) (McLellan *et al*, 2011) RSVF.

### Preparation for dose-response analysis

Peptide reports generated in Spectronaut were processed using an in-house R script in R Statistical Software (version 4.2.0; R Core Team 2021). In brief, proteins with at least two peptide precursors were considered. RSVF and AC8-CaM datasets included only proteotypic peptides. For aSyn datasets, both proteotypic and non-proteotypic peptides were covered; therefore, non-proteotypic peptides, which are reported, should be taken with caution. Raw peptide abundances were normalized using variance stabilizing normalization with the *vsn* package (Huber *et al*, 2002) (version 3.64.0). We used an outlier detection method based on the interquartile range (IQR) to define boundaries outside of the first (Q1) and third (Q3) quartile for peptide abundances within each condition per peptide precursor. Peptide abundances that are more than 1.5 times the IQR below Q1 or more than 1.5 times above Q3 are considered outliers. This approach ensured that potential outliers are removed prior to dose-response analysis. Subsequently, we considered only peptide precursors that were measured in at least 3 replicates, covering at least 5 conditions. Filtered data, consisting of normalized log_2_-transformed peptide abundances, were scaled to a range between 0 and 1 and subjected to dose-response analysis using the *protti* package (Quast *et al*, 2022) (version 0.5.0) that utilizes the log-logistic model with four parameters (LL.4) from the *drc* package (Ritz *et al*, 2015) (version 3.0-1). Pearson’s correlation coefficient *r* was used to assess the strength of the sigmoidal trend of dose-response profiles. Only peptides that fulfilled the Benjamini-Hochberg-corrected *p*-values (*q*-values) < 0.01 obtained from an analysis of variance (ANOVA) and Pearson’s correlation coefficients *r* > 0.85 were considered significant. Unscaled peptide abundances were used for statistical testing of differentially abundant peptides using an empirical Bayes moderated t-test as implemented in the *limma* package (Ritchie *et al*, 2015) (version 3.52.4). The resulting *p*-values were adjusted by multiple testing using the Benjamin-Hochberg method. The output of the statistical analysis was filtered using the following cutoffs: *q*-value < 0.01 and |log_2_ FC| > 0.75. The final list of peptides that fulfilled all the defined criteria represented differentially altered peptides. Every differentially altered peptide has an EC50 value assigned, which represents the inferred quantity of a protein necessary to observe half-maximum of the relative peptide intensity change between treated and untreated samples.

### Proteomic analysis of CRISPR/Cas9-mediated knockout of SNCA in iPSC-derived cortical neurons

To verify the knockout of SNCA, we analyzed the expression levels of aSyn in both healthy control (n = 4) and SNCA-KO (n = 4) iPSC lines using quantitative DIA–MS. MS2 quantification of aSyn peptides was performed with Spectronaut as described above. Extracted ion chromatographs for peptides of aSyn (Figure S 2e) were exported from Spectronaut 15.

### Mapping of known interactors

Systematic analysis of known interactors was conducted using the STRING database (https://string-db.org) of physically interacting proteins (Szklarczyk *et al*, 2021). Proteins with a score of >150 were considered known interactors. Fisher’s exact test (*p*-value < 0.01) was used to determine whether known interactors are enriched amongst our identified proteins relative to all identified known interactors.

### Identification of Lewy body (LB)-associated proteins

To evaluate whether structurally altered proteins upon spike-in of aSyn are associated with the formation of LBs, we utilized data from previous studies which report on LB-associated proteins in a neuronal aSyn fibril seeding model (Mahul-Mellier *et al*, 2020) or in post-mortem patient brains (Petyuk *et al*, 2021; Xia *et al*, 2008) Fisher’s exact test (*p*-value < 0.01) was used to determine whether LB-associated proteins are enriched amongst our identified structurally altered proteins relative to all identified LB-associated proteins. The web-based functionality g:Orth was used for orthology search to translate *M. musculus* genes into *H. sapiens* genes using g:Profiler (version e106_eg53_p16_65fcd97, database updated on 18/05/2022) (Raudvere *et al*, 2019).

### Analysis of calmodulin-binding motifs

Calmodulin (CaM)-binding motifs were assessed using an in-house R script in R Statistical Software (version 4.2.0; R Core Team 2021). Briefly, we concatenated information about known CaM-binding motifs from the Calmodulation database and Meta-analysis predictor website (http://cam.umassmed.edu), which enables the prediction of CaM-binding motifs in protein sequences (Mruk *et al*, 2014). We predicted the presence of CaM-binding motifs in structurally altered peptides and calculated the number of CaM-binding motifs per protein (without discrimination for transmembrane domains).

### Functional enrichment analysis

Functional enrichment analysis of proteins, based on gene ontology (GO) terms molecular function (MF), biological process (BF), and cellular component (CC), was performed using g:Profiler (version e106_eg53_p16_65fcd97, database updated on 18/05/2022) (Raudvere *et al*, 2019) with the FDR multiple testing correction method applying a significant threshold of 0.01 (*q*-value < 0.01). We further utilized the *rrvgo* package (Supek *et al*, 2011) (version 1.8.0) to summarize the enriched terms by removing redundant GO terms. The resulting list size was set to 0.7 and the SimRel functional similarity measure for comparing two GO terms with each other was considered.

### Network propagation and clustering analysis

To assess associations between our identified proteins and PD-related traits in the aSyn experiment, we performed network-based expansion and clustering analysis as described previously (Barrio-Hernandez *et al*, 2021). In brief, to generate a list of starting genes for network expansion, we used all proteins from the LiP experiments and all genes linked to different PD-related traits. We defined a list of Parkinson-related disorders based on the EFO hierarchy, selecting all traits that have the term “Parkinson’s disease (EFO:0002508)” as the ancestor and genes with associations based on common or rare variants, leading to the following list: “Parkinson disease, mitochondrial”, “Young adult-onset Parkinsonism”, “Parkinson’s disease” and “Hereditary late-onset Parkinson’s disease”. To select genes associated with a given trait, we used the evidence present in the OpenTargets platform (https://www.opentargets.org/). For common variants, we selected all genes with an L2G score (association of an SNP to a given gene from GWAS studies) bigger than 0.5. For rare variants, we used all genes linked to SNPs with a clinical output not considered “benign”, according to ClinVar definitions (https://www.ncbi.nlm.nih.gov/clinvar/). The network expansion is performed trait-per-trait, following an approach described previously (Barrio-Hernandez *et al*, 2021). Briefly, we first mapped all the genes considered as starting signals (LiP experiments or genes associated with genetic evidence of association to a given disease) to a custom version of OpenTargets interactome (compilation of interactions from IntAct, Reactome, Signor, and STRING with score >= 0.75). We applied network propagation using the personalized PageRank algorithm included in the *igraph* package (Csárdi & Nepusz, 2006) (version 1.2.4.2). Those genes with a ranking score bigger than the Q3 (75% of the distribution) are selected for community detection using the walktrap algorithm from the *igraph* package (version 1.2.4.2) via random walks. To define significant communities, we compared the PageRank score, resulting from the network propagation inside and outside the community, using the Kolmogorov-Smirnov test with the Benjamini-Hochberg adjustment. We selected the communities with at least 1 starting hit, no less than 10 nodes in total, and with an adjusted *p*-value smaller than 0.05. Significant communities were compared among traits by measuring the nodes’ overlap using the Jaccard index. To calculate an enrichment based on GOBP annotation, Fisher’s exact test (*p*-value < 0.05) was used.

### Data analysis of enriched and depleted proteins

To identify proteins that are either enriched in the pellet with aSyn amyloid fibrils or depleted from the supernatant, log_2_-transformed protein abundances provided in the Spectronaut report were used to calculate log_2_ FC and *q*-values using an empirical Bayes moderated t-test as implemented in the *limma* package (Ritchie *et al*, 2015) (version 3.52.4). For the assessment of pellet-enriched proteins, pellet samples containing aSyn amyloid fibrils were compared to samples treated with aSyn monomer or untreated samples. For supernatant-depleted proteins, the supernatant recovered after ultracentrifugation was used. Significant proteins (*q*-value < 0.01) that changed in abundance either in the pellet or supernatant samples with a fold-change of 1.5 were considered enriched or depleted. Fisher’s exact test (*p*-value < 0.01) was used to determine whether putative fibril-binding proteins obtained by ultracentrifugation are enriched amongst our identified structurally altered proteins relative to all identified proteins.

### Prediction of AC8 structure

The 3D structure of bovine AC8 was predicted from its amino acid sequence using AlphaFold2 (Varadi *et al*, 2022; Jumper *et al*, 2021). The FASTA file containing the amino acid sequence of AC8 and its tags was submitted to the AlphaFold prediction algorithm. The predicted structure was then used to visualize differential peptides upon CaM treatment.

### 3D analysis of protein structural changes

Significantly altered peptides were mapped onto representative 3D protein structures obtained from the Protein Data Bank (Berman *et al*, 2000). In RSVF experiments, preRSVF (PDB: 4JHW) (McLellan *et al*, 2013) and/or postRSVF (PDB: 3RRR) (McLellan *et al*, 2011) structures were used to identify RSVF regions that changed due to antibody binding. In AC8-CaM experiments, the predicted structure (uploaded to the public repository) was used to detect CaM-binding sites of AC8. All structural alterations were visualized using the molecular visualization program UCSF ChimeraX (1.3rc202111292147) (Goddard *et al*, 2018; Pettersen *et al*, 2021).

## Data Availability

The mass spectrometry proteomics data have been deposited to the ProteomeXchange Consortium via the PRIDE partner repository (Perez-Riverol *et al*, 2022) with the dataset identifiers PXD039481, PXD039520, and PXD039784.

## Code Availability

The custom R scripts developed and used in this study are available via GitHub at https://gitfront.io/r/PicottiGroup/FeTezEanyUFM/LiP-MS-protein-protein-interactions-lip-data-structural-analysis-protein-protein-interactions/.

## Acknowledgments

The authors thank the Correia laboratory (EPFL Lausanne, Switzerland) for providing samples of purified RSVF and site-specific antibodies for validation experiments. P.P. was supported by the European Research Council (grant agreement no. 866004), the EPIC-XS Consortium (grant agreement no. 823839), a Sinergia grant from the Swiss National Science Foundation (SNSF grant CRSII5_177195), and grants from Parkinson Schweiz and the Empiris Foundation. V.M.K. was supported by a Swiss National Science Foundation grant (no. 184951). W.H. was supported by the Oxford-McGill-Zurich Partnership in Neuroscience and is currently supported by UZH URPP AdaBD. P.B. is supported by the Helmut Horten Stiftung and the ETH Zurich Foundation.

## Ethics approval and consent to participate

Derivation of the human iPSC line SFC856-03-04 (used as the starting point for SNCA knockout) used in this study is described elsewhere (Haenseler *et al*, 2017b). The iPSC line was derived from dermal fibroblasts from healthy donors through the Oxford Parkinson’s Disease Centre: participants recruited to this study had signed informed consent, which included the derivation of human iPSC lines from skin biopsies (Ethics Committee that specifically approved this part of the study: for control donors, National Health Service, Health Research Authority, NRES Committee South Central, Berkshire, UK, REC 10/H0505/71, and for SNCA patient REC 07/H0720/161).

## Author contributions

A.H. and P.P. conceived the experimental design of the study. A.H. performed the experiments, performed mass spectrometric measurements of the RSVF and aSyn datasets, carried out data analysis and interpretation of all datasets presented in this study, and wrote the manuscript with N.d.S. and with contributions from P.P. and from all authors. D.S. designed and performed the experiments of the AC8-Calmodulin dataset, and carried out mass spectrometric measurements. F.S. provided samples of purified RSVF and antibodies for all RSVF experiments. P.S. performed quality control experiments of aSyn. J.V. and S.A.C. designed and performed the knockout of SNCA in iPSC-derived cortical neurons. W.H. provided pellets of iPSC-derived cortical neurons. I.B-H. carried out network-based expansion and clustering analysis. D.G. purified the N-terminally acetylated aSyn and performed quality control experiments. L.N. developed the script to predict CaM-binding domains via canonical CaM-binding motifs. P.P. conceived and supervised the study.

